# Multi-kinesin clusters impart mechanical stress that reveals mechanisms of microtubule breakage in cells

**DOI:** 10.1101/2025.01.31.635950

**Authors:** Qi Geng, Andres Bonilla, Siara N. Sandwith, Kristen J. Verhey

**Affiliations:** Department of Molecular, Cellular, and Developmental Biology, University of Michigan, Ann Arbor, MI, USA; Department of Cell and Developmental Biology, University of Michigan Medical School, Ann Arbor, MI, USA; Cellular and Molecular Biology Program, University of Michigan Medical School, Ann Arbor, MI, USA; Program in Biophysics, University of Michigan, Ann Arbor, MI, USA

## Abstract

Microtubules are cytoskeletal filaments that provide structural support for numerous cellular processes. Despite their high rigidity, microtubules can be dramatically bent in cells and it is unknown how much force a microtubule can withstand before breaking. We find that liquid-liquid phase separation of the kinesin-3 motor KIF1C results in multi-kinesin clusters that entangle neighboring microtubules and impose a high level of mechanical stress that results in microtubule breakage and disassembly. Combining computational simulations and experiments, we show that microtubule fragmentation is enhanced by having a highly processive kinesin motor domain, a stiff clustering mechanism, and sufficient drag force on the microtubules. We estimate a rupture force for microtubules in cells of 70-120 pN, which is lower than previous estimates based on *in vitro* studies with taxol-stabilized microtubules. These results indicate that the presence of multiple kinesins on a cargo has the potential to cause microtubule breakage. We propose that mechanisms exist to protect microtubule integrity by releasing either the motor-cargo or motor-microtubule interaction, thereby preventing the accumulation of mechanical stress upon the engagement of multi-motor clusters with microtubules.

## Introduction

Microtubules are a major component of the eukaryotic cytoskeleton and provide structural and mechanical support for a variety of fundamental cellular processes, including intracellular trafficking, beating of cilia, cell division, and cell migration. Microtubules are self-assembled via the head-to-tail (longitudinal) binding of α/β-tubulin subunits into polarized protofilaments, and via lateral interactions of tubulin subunits that wrap ∼13 parallel protofilaments into a hollow tube. The large number of interactions between tubulin subunits make microtubules the stiffest of the cytoskeletal filaments, with a rigidity comparable to Plexiglas (Gittes et al., 1993). This high stiffness allows microtubules to sustain mechanical stress during processes that generate substantial force such as intracellular transport, the positioning of nuclei during cell migration, and the separation of chromosomes during cell division (Guo et al., 2013; Zheng et al., 2020; Risteski et al., 2021). However, it is still unclear how much force a microtubule can sustain before breaking.

As a comparison, the material strength of actin filaments (F-actin) is better characterized. Manipulating a single actin filament by microneedles *in vitro* measured the actin-actin bond breaking force to be ∼600 pN for an untwisted phalloidin-stabilized filament, which decreases to as low as ∼200 pN when filaments are under torsion (Tsuda et al., 1996). It is assumed that microtubules require a higher breaking force based on the larger number of inter-subunit H-bonds for tubulin subunits in a microtubule filament (∼8 longitudinal H-bonds per protofilament x 13 protofilaments = ∼104 longitudinal H-bonds) than for actin subunits in F-actin (∼18 H-bonds) (Endow and Marszalek, 2019; Merino et al., 2018; Zhang et al., 2018). Consistent with this, a fully-atomistic molecular dynamics simulation found that microtubules can sustain ∼150 MPa of tensile stress (stretching), which corresponds to a force of ∼55,000 pN (Wu and Adnan, 2018).

However, experimental studies have suggested that microtubules subjected to stress applied along their longitudinal axis break when subjected to much lower forces. Using a stretchable polydimethylsiloxane (PDMS) substrate to exert a tensile force to taxol-stabilized microtubules, Kabir et al. reported a minimal longitudinal strain of 4.28% for breaking microtubules (Kabir et al., 2014). With Young’s modulus estimated to be ∼12 MPa in this paper, a breaking force of ∼170 pN can be derived. Using Young’s modulus of ∼1.2 GPa (Gittes et al., 1993; Kurachi et al., 1995) gives a microtubule rupture force of

∼17,000 pN. Interestingly, less microtubule breaking was observed upon application of compressive forces using a similar experimental setup and required experimental conditions (e.g. high concentrations of anchors or microtubule pinning) that generate tight buckling (low radii of curvature) (Kabir et al., 2015, 2020; Tsitkov et al., 2022; Nasrin et al., 2024). These results suggest that microtubules are more sensitive to tensile forces than to compressive forces.

Another study observed microtubule breaking upon longitudinal stretching and compression in a gliding assay with a mutant kinesin-14 motor that walks towards both plus and minus ends of microtubules. In this assay, taxol-stabilized microtubules were observed to undergo abrupt reversals of direction, buckling events, and breaking events due to motor pulling and pushing forces. Estimating the number of active motors pulling on individual microtubules suggested a minimum rupture force of ∼500 pN (Endow and Marszalek, 2019). The estimated breaking force for individual microtubules thus differs by several orders of magnitude.

Furthermore, it is unclear how these studies, which utilized taxol-stabilized microtubules, can be interpreted with respect to microtubules in cells. Indeed, live-cell imaging has revealed that microtubules undergo extensive bending and buckling in cells, yet they rarely break or fragment (Waterman-Storer and Salmon, 1997; Odde et al., 1999; Wang et al., 2001; Gupton et al., 2002; Brangwynne et al., 2006; Bicek et al., 2009). The underlying processes that lead to microtubule bending and buckling have been suggested to include their polymerization against the cell cortex, their transport or sliding by kinesin and dynein motor proteins, and contraction of the surrounding actomyosin system (Schaefer et al., 2002; Bicek et al., 2009). Of these, it seems that only actomyosin contraction is capable of producing sufficient force for microtubules to rupture as microtubule breakage is more frequently observed at the leading edge of motile cells (Waterman-Storer and Salmon, 1997; Gupton et al., 2002). Although the rupture force, a fundamental microtubule feature, has not been directly measured in cells, it was estimated that microtubules can withstand ∼100 pN compressive force before buckling in cells (Brangwynne et al., 2006). The ability of individual microtubules to bear such large compressive forces is thought to be due to lateral mechanical reinforcement by the surrounding elastic cytoskeleton (Brangwynne et al., 2006; Li et al., 2023; Ju et al., 2024; Orii and Tanimoto, 2024).

Our previous study showed that the kinesin-3 motor KIF1C undergoes liquid-liquid phase separation (LLPS) through an intrinsically disordered region (IDR) in its C-terminal tail domain (Geng et al., 2024). LLPS results in membrane-less condensates localized at the cell periphery and enriched with KIF1C proteins and associated mRNA molecules. Live-cell imaging demonstrated that the condensates are highly motile (Geng et al., 2024), suggesting that when KIF1C proteins are concentrated in the condensates, their motor domains can still interact with and move along microtubules.

Here, we report that overexpression of the kinesin-3 motor KIF1C in cultured cells results in multi-motor clusters that entangle neighboring microtubules and cause them to bend, buckle, and break. Although microtubule breaking in this system is an artifact of KIF1C overexpression, it provides an experimental setup to probe microtubule breakage in the intracellular environment. We used Cytosim for computational modeling (Nedelec and Foethke, 2007; Lugo et al., 2023) of microtubule breaking behavior by simulating the interaction between multi-kinesin clusters and a network of microtubules. We find that microtubule breakage is greater under conditions where microtubules experience high drag force from cytoplasmic viscosity and/or anchoring proteins. The simulations also predict that the level of microtubule fragmentation depends on i) the biophysical properties of individual motors in the clusters, such as stall force, velocity, and processivity and ii) properties of the clustering mechanism, such as the stiffness of cluster and the number of motors in a cluster. These predictions were tested and verified experimentally in cells, giving rise to an estimated force of 70-120 pN required to break microtubules in cells. The limited amount of microtubule breakage observed under normal cellular conditions suggests that there must be mechanisms to prevent microtubule fragmentation. We hypothesized that one mechanism that can prevent the buildup of mechanical stress that leads to microtubule breakage is a labile kinesin-cargo interaction and indeed, we find that clustering of kinesins via a protein-lipid interaction can prevent microtubule breakage in cells.

## Results

### Bimolecular condensates of KIF1C induce microtubule fragmentation in cells

To gain insight into the interaction of KIF1C in condensates with microtubule tracks, we fixed COS-7 cells expressing fluorescently-tagged KIF1C and labeled the microtubules by immunofluorescence. Strikingly, numerous short microtubule fragments were observed near the KIF1C condensates (Fig. 1 A). This phenomenon could be observed in other cell types such as hTERT-RPE1 cells (Fig. 1 A). The number of microtubule fragments in cells expressing KIF1C-GFP positively correlates with KIF1C’s LLPS activity (Fig. 1 B). We noticed that KIF1C condensates colocalize with what appear to be entangled and highly dense areas of microtubules, while the already fragmented pieces of microtubules primarily localize peripheral to KIF1C condensates (Fig. 1 A). This suggests that the microtubule fragments may be produced from a process involving KIF1C condensates interacting with a collection of microtubules.

**Figure 1.**
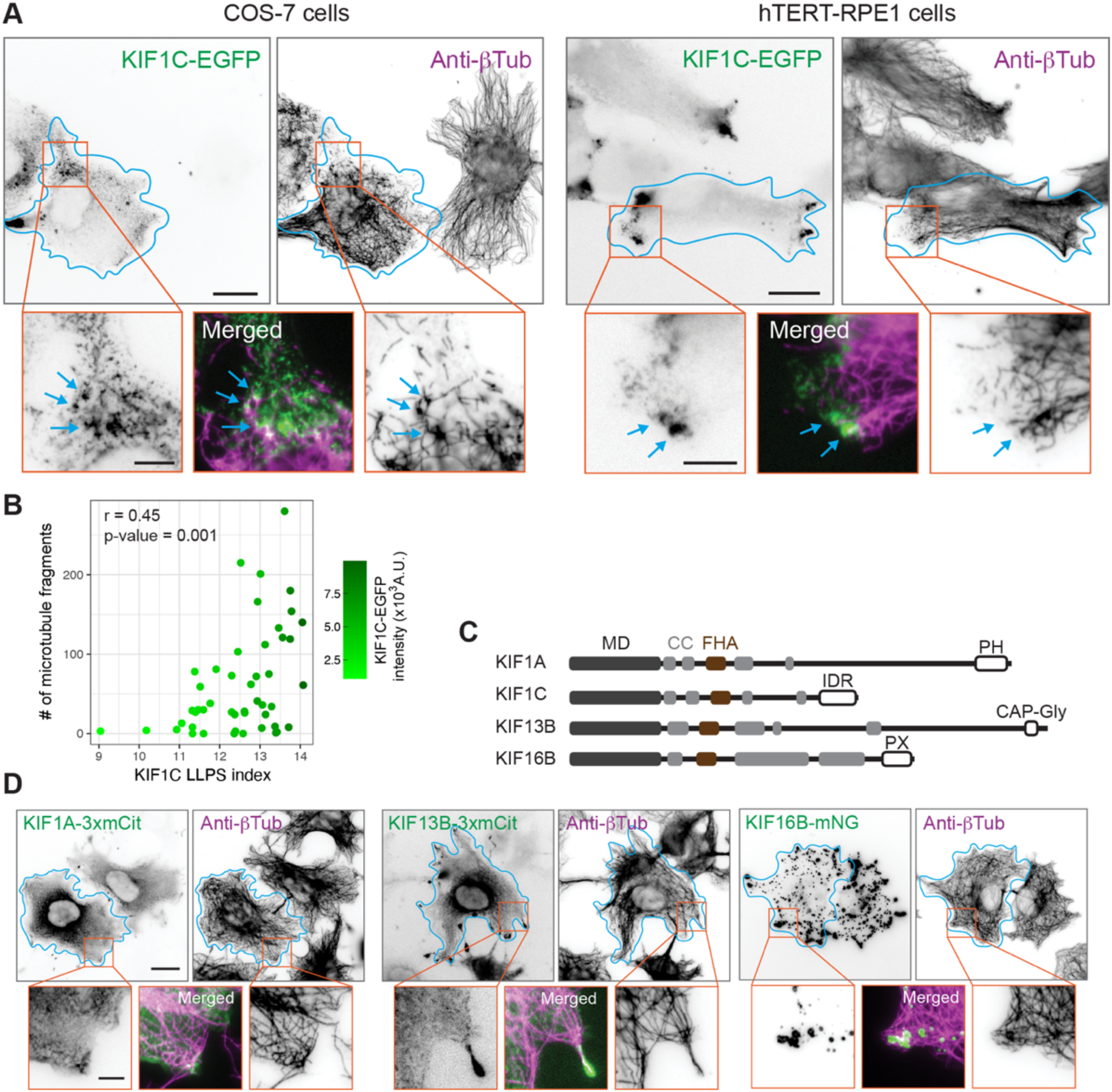
KIF1C biomolecular condensates induce MT fragmentation in cells. **(A)** Representative images of COS-7 and hTERT-RPE1 cells expressing KIF1C-EGFP and fixed and stained for microtubules (anti-β-tubulin). Cyan arrows point to colocalization of KIF1C-EGFP condensates and dense microtubules. **(B)** Correlation between KIF1C phase separation and MT fragmentation quantified in COS-7 cells. X-axis: LLPS index. Y-axis: number of MT fragments. Each dot represents one cell (N = 48 cells from one representative experiment). The color of each dot reflects the total intensity of KIF1C-EGFP fluorescence in that cell. The Pearson correlation coefficient (r) and associated p-value correlating KIF1C-EGFP intensity heterogeneity and the number of MT fragments are shown in the plot. **(C)** Schematic of domain organization of the indicated kinesin-3 family members. MD: Motor domain; CC: Coiled-coil domain; FHA: Forkhead-associated domain. PH: Pleckstrin homology domain; IDR: Intrinsically disordered region; CAP-Gly: Glycine-rich domain of Cytoskeleton-associated proteins; PX: PhoX homology domain. **(D)** Representative images of COS-7 cells expressing fluorescently-tagged KIF1A, KIF13B or KIF16B fixed and stained for β-tubulin to show the microtubule distribution. For all microscopy images of this figure, cyan lines mark the cell outline and orange boxes indicate the regions shown in the magnified images. Scale bars: 20 μm for whole-cell views and 5 μm for magnified images.

As a control, we transfected COS-7 cells with fluorescently-tagged constructs of other kinesin-3 motors which share a conserved N-terminal motor domain with KIF1C but have different C-terminal tail domains (Fig. 1 C). None of the other kinesin-3 motors produced microtubule fragments when expressed under the same conditions (Fig. 1 D), indicating that the appearance of microtubule fragments specifically correlates with KIF1C expression. As an additional control, we generated a rigor mutant of KIF1C by mutating a glycine to alanine (G251A) in the conserved switch II motif (Kikkawa et al., 2001). The rigor mutant KIF1C(G251A)-mNG showed strong microtubule decoration but did not cause microtubule fragmentation (Fig. S1), indicating that the appearance of microtubule fragments requires the motility of KIF1C along microtubules.

The distinguishing feature of KIF1C within the kinesin-3 family is its IDR that drives LLPS (Fig. 1 C). To test whether phase separation is sufficient to produce microtubules fragments, we expressed a truncated version of KIF1C containing only the stalk and tail regions (ST, aa 349-1103), which forms condensates in cells (Geng et al., 2024). However, KIF1C(ST) condensates did not cause the appearance of fragmented microtubules in cells (Fig. 2 A). We thus tested whether the motor domain (MD, aa 1-376) is sufficient for generating microtubule fragments, as has been shown previously for the kinesin-1 KIF5 motor domain (Triclin et al., 2021; Andreu-Carbó et al., 2022; Budaitis et al., 2022). The KIF1C(MD) protein showed peripheral accumulation with clear microtubule decoration, a signature of highly processive kinesin-3 motor domains (Soppina et al., 2014), but it did not cause the appearance of fragmented microtubules (Fig. 2 A). These results suggest that neither the LLPS activity nor the microtubule-based motility of KIF1C are sufficient to generate microtubule fragmentation in cells.

**Figure 2.**
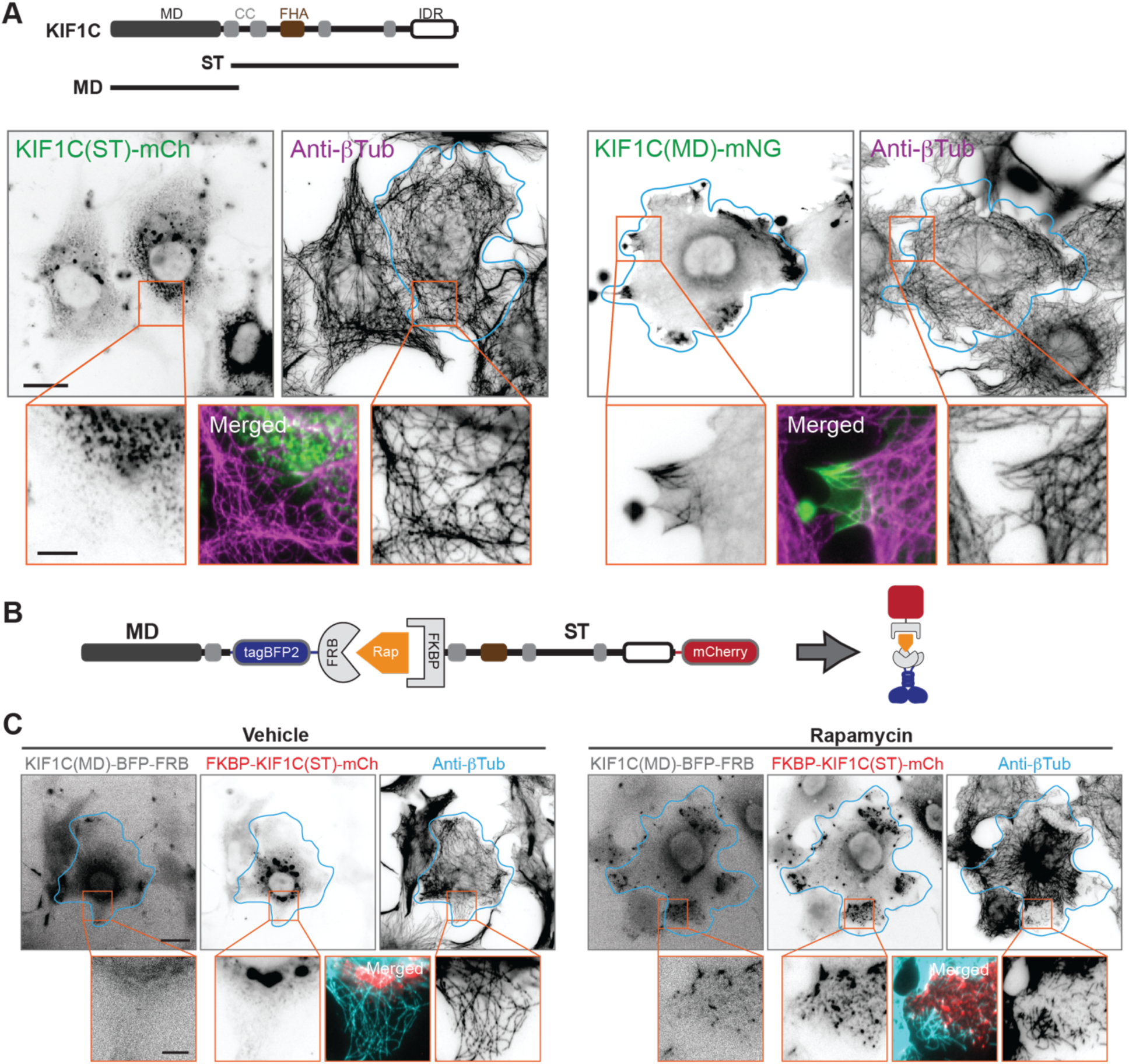
KIF1C motility and LLPS activity are required for MT fragmentation. **(A)** Top: Schematic of full-length KIF1C (aa 1-1103) and the truncations stalk+tail (ST, aa 349-1103) and motor domain (MD, aa 1-376). Bottom: Representative images of COS-7 cells expressing fluorescently-tagged KIF1C(ST) and KIF1C(MD) constructs fixed and stained for microtubules (anti-β-tubulin). **(B)** Schematic of the split-kinesin assay. The kinesin motor domain is fused to tagBFP2 and FRB whereas the kinesin stail+tail domains are fused to mCherry and FKBP. Addition of rapamycin results in heterodimerization of FRB and FKBP and reformation of a full-length kinesin. **(C)** Representative images of COS-7 cells expressing KIF1C split-kinesin components and fixed and stained for microtubules (anti-β-tubulin). Left, cells treated with vehicle control (ethanol); right, cells treated with 44 nM rapamycin for 30 min before fixation. Cyan lines mark the cell outline, orange boxes indicate the regions shown in the magnified images. Scale bars: 20 μm for whole-cell views and 5 μm for magnified images.

### A split-kinesin assay reveals the breakage of microtubules by KIF1C in real time

To verify that microtubule fragmentation requires both LLPS and motility of KIF1C, we utilized the chemically-induced split-kinesin assay (Muthuswamy et al., 1999; Jenkins et al., 2012). This assay enables us to combine the LLPS behavior of the KIF1C(ST) and the motility behavior of the KIF1C(MD) in a temporally-controlled manner in cells (Fig. 2 B). In the absence of rapamycin (vehicle control), KIF1C(MD)-tagBFP2-FRB accumulated at the cell periphery and FKBP-KIF1C(ST)-mCherry localized in cytoplasmic puncta and the microtubules did not appear broken or fragmented, as expected (Fig. 2 C). In contrast, upon treatment with 44 nM rapamycin for 30 min, KIF1C(MD)-tagBFP2-FRB and FKBP-KIF1C(ST)-mCherry colocalized in condensates at the cell periphery and were surrounded by entangled and/or fragmented microtubules (Fig. 2 C). These results indicate that both the LLPS of KIF1C and its motility along microtubules are required to generate microtubule fragments in cells.

We thus used the split-kinesin assay and live-cell imaging to capture the generation of microtubule fragments in real time. In live COS-7 cells, KIF1C(MD) + KIF1C(ST) condensates were assembled upon rapamycin treatment and were observed to move along microtubule tracks towards the cell periphery, often switching tracks or pausing at microtubule intersections (Video 1 and 2; Fig. 3 A,B and Fig. S2). Upon reaching the cell periphery, the condensates remained associated with the plus ends of the microtubules and underwent fusion and fission events to form larger or smaller structures. They also caused extensive sliding, bending, and twisting of microtubules and despite the density of the microtubule network, microtubule breaking events could be observed. The newly generated microtubule fragments moved away from the entangled microtubule network and became more visible (Fig. 3 B top, Fig S2). On occasion, the KIF1C condensates engaged microtubules in the central region of the cell in a manner that caused them to break (Fig. 3 B bottom). These events were relatively rare, presumably because the central microtubule network is anchored in place.

**Figure 3.**
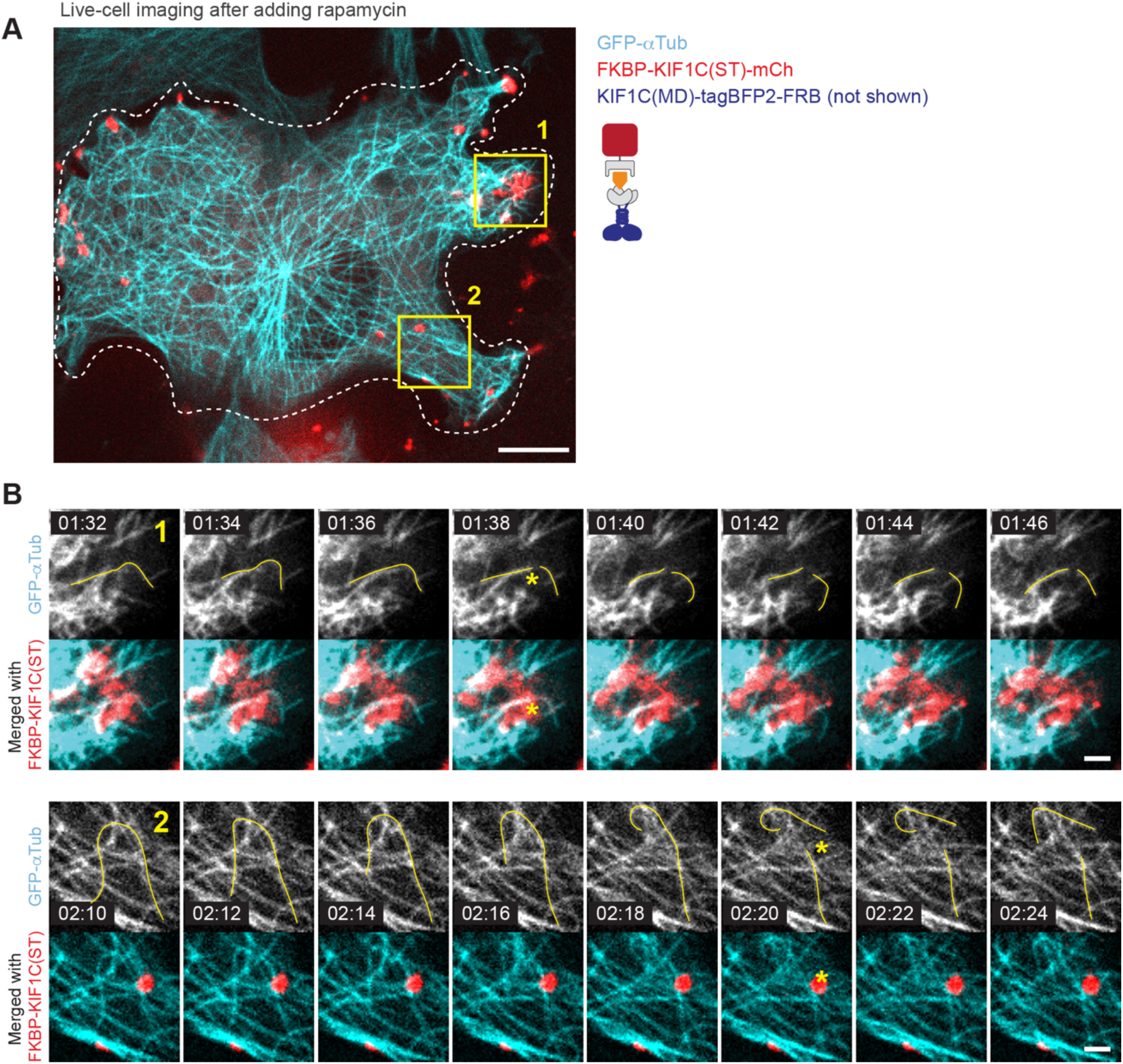
Live imaging of MT breakage by KIF1C condensates. **(A)** Representative example of live-cell imaging of COS-7 cells expressing KIF1C(MD)-tagBFP2-FRB (imaging channel not shown), FKBP-KIF1C(ST)-mCherry (magenta), and EGFP-α-tubulin (cyan). The cells were imaged at 2 frames per second starting 10-15 min after rapamycin addition. Dashed white lines mark the cell outline, yellow boxes indicate the regions magnified in panel (B). Scale bar: 10 μm. **(B)** Time lapse images of microtubule breaking in the boxed regions in (A). Top, grayscale images of the EGFP-α-tubulin channel. Bottom, merge of the FKBP-KIF1C(ST)-mCh (magenta) and EGFP-α-tubulin (cyan) channels. The yellow lines indicate condensate-associated MTs that broke during the time course. The yellow asterisks mark the time-frame (min:sec) and position of MT breaking. Scale bar, 2 μm.

Based on these observations, we propose that the microtubules are fragmented by mechanical forces generated by KIF1C motors in the condensates. Specifically, KIF1C motor proteins that self-assemble into a biomolecular condensate by phase separation can simultaneously interact with multiple neighboring microtubules and can entangle the microtubules by sliding and crosslinking them. The mechanical stress that accumulates in the system results in the breaking of microtubules (Fig. 4 A).

**Figure 4.**
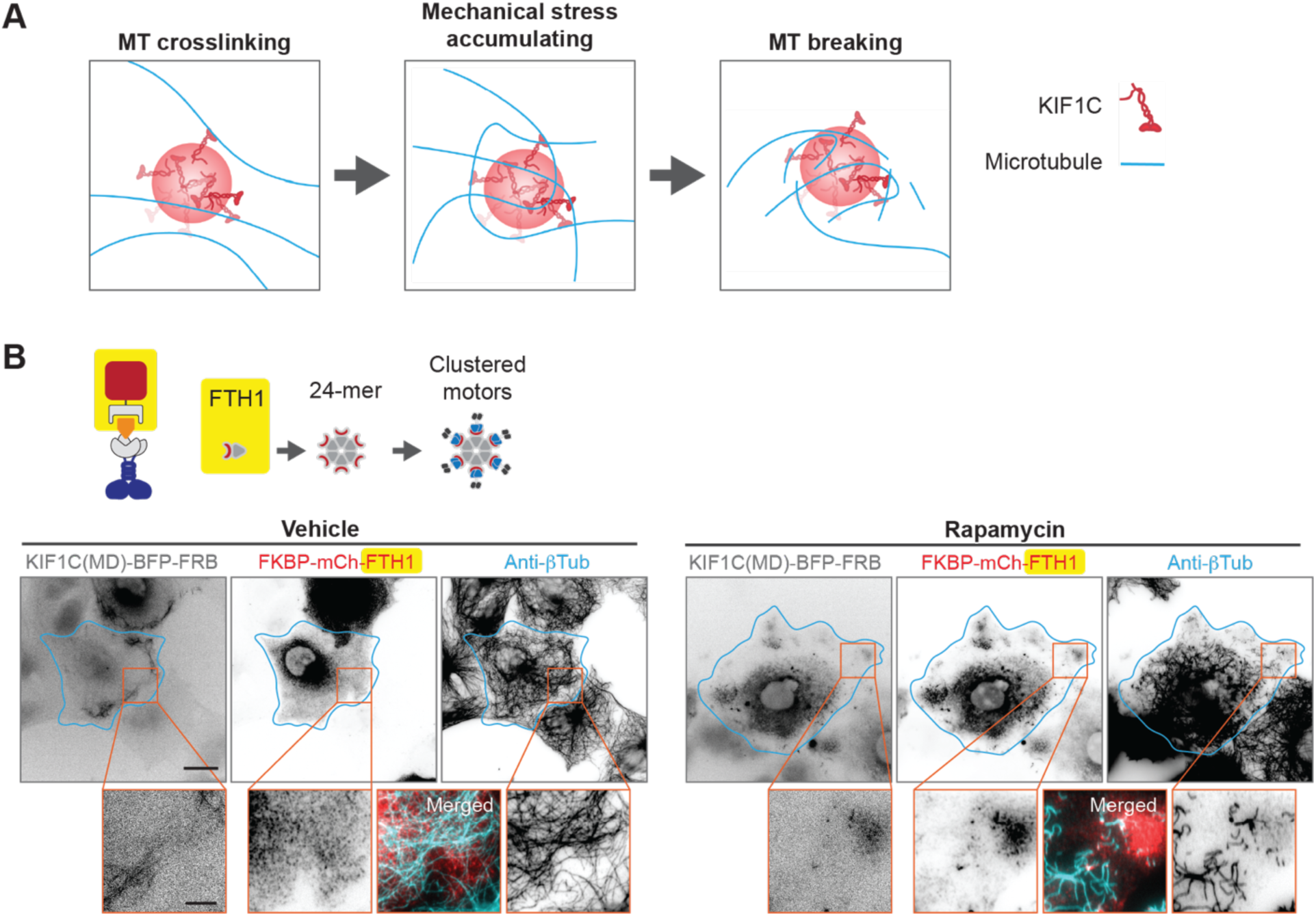
Clustering of kinesin motor domains is key to KIF1C-induced microtubule fragmentation. **(A)** Proposed model of how the clustering of KIF1C proteins into phase-separated condensates leads to microtubule breakage and fragmentation. **(B)** Representative images of COS-7 cells expressing KIF1C(MD)-tagBFP2-FRB and FKBP-mCh-FTH1 and fixed and stained for microtubules (anti-β-tubulin). Left, cells treated with vehicle control (ethanol); right, cells treated with 44 nM rapamycin for 30 min before fixation. Cyan lines mark the cell outline, orange boxes indicate the regions shown in the magnified images. Scale bars: 20 μm for whole-cell views and 5 μm for magnified images.

A key feature of this model is the self-assembly of multiple kinesins into a molecular cluster in a manner that maintains their ability to interact with and move along microtubules. We thus reasoned that other mechanisms of generating such multi-KIF1C clusters should be able to recapitulate microtubule fragmentation. To test this, we replaced the KIF1C(ST) construct in the split-kinesin assay with the human ferritin heavy chain (FTH1) protein known to self-assemble into a 24-mer spherical particle (Lawson et al., 1991; Bracha et al., 2018). FTH1 was tagged with FKBP and mCherry (FKBP-mCherry-FTH1) for recruitment of FRB-tagged KIF1C motor domains in a temporally-controlled manner. In the absence of rapamycin, the KIF1C(MD) accumulated at the plus ends of microtubules in the cell periphery, the FTH1 protein was largely cytoplasmic, and the microtubules appeared intact. Upon rapamycin treatment, the assembled KIF1C(MD) + FTH1 particles colocalized at the cell periphery and microtubule fragmentation could be clearly observed (Fig. 4 B).

### Computational modeling of microtubule breaking by multi-kinesin clusters

Although microtubule breakage is an artifact of overexpression of a protein that self-assembles and undergoes microtubule-based motility, we reasoned that it could be used to gain a quantitative understanding of how microtubules respond to mechanical stress. However, it is not known how much force multiple KIF1C proteins in a cluster can produce. Nor is it known how much force is required to break microtubules in cells. Given these limitations, we utilized Cytosim, a cytoskeleton simulation tool that has been used previously for mesoscale modeling of microtubule-motor interactions (Nedelec et al., 1997; Surrey et al., 2001; Nedelec and Foethke, 2007; Lugo et al., 2023), for computational modeling of the interaction between multi-kinesin clusters and the microtubule network.

We utilized the existing functionality of Cytosim to configure cluster objects with a defined number of motors on their surface and fiber objects to mimic the microtubule network. As Cytosim does not allow microtubule breakage, we first modified the source code to enable the breaking of microtubule fibers upon mechanical stress. The code evaluates the tension experienced by each microtubule fiber at each time step of the simulation and then implements a breakage event based on a threshold force for microtubule breaking that can be defined in the configuration file.

We tested and validated this microtubule fiber breaking functionality at the level of single microtubules. We set up simulations in which artificial motors (Fig. S3 A, green circles) holding the ends of a microtubule fiber walk away from each other and thereby apply a stretching force to the microtubule. The maximum tension that can be applied to the microtubule fiber depends on the stall force (F_stall_) of the artificial motors. As expected, when the threshold force for microtubule breaking was set to a value equal to F_stall_, the fiber readily snapped into two fragments at the position experiencing the highest local tension (Fig. S3 B). Microtubule breaking was prevented only under conditions in which the microtubule breaking threshold was set at a value higher than maximum tension that can be applied to the microtubule fiber (Fig. S3 C).

We then created a simulation space of 20 μm x 20 μm to mimic an area at the periphery of a cell. A total of 60 microtubule fibers were randomly placed into the space to form a microtubule network with frequent intersections. Multi-motor clusters containing 24 motors on their surface, mimicking the FTH1-mediated kinesin clustering (Fig. 4 B), were also distributed randomly into the space as were components that are likely important for the organization of the microtubule network such as crosslinkers that attach microtubules to each other and anchors that attach microtubules to the space (Fig. 5 A). The default properties of the motor, including stall force, velocity, and processivity were set to those of KIF1C MD, as measured in previous work (Siddiqui et al., 2022) or this study (Fig. S4 A,B). Microtubule rigidity and cytoplasmic viscosity were set based on measurements in the literature (Gittes et al., 1993). With the goal of identifying conditions that produce microtubule fragmentation similar to that observed in live cells (Fig. 3), we tested a range of values for each parameter to arrive at default values (Table 1). Finally, to determine how much force microtubules can withstand before breaking, we tested a range of threshold values for microtubule breaking.

**Figure 5.**
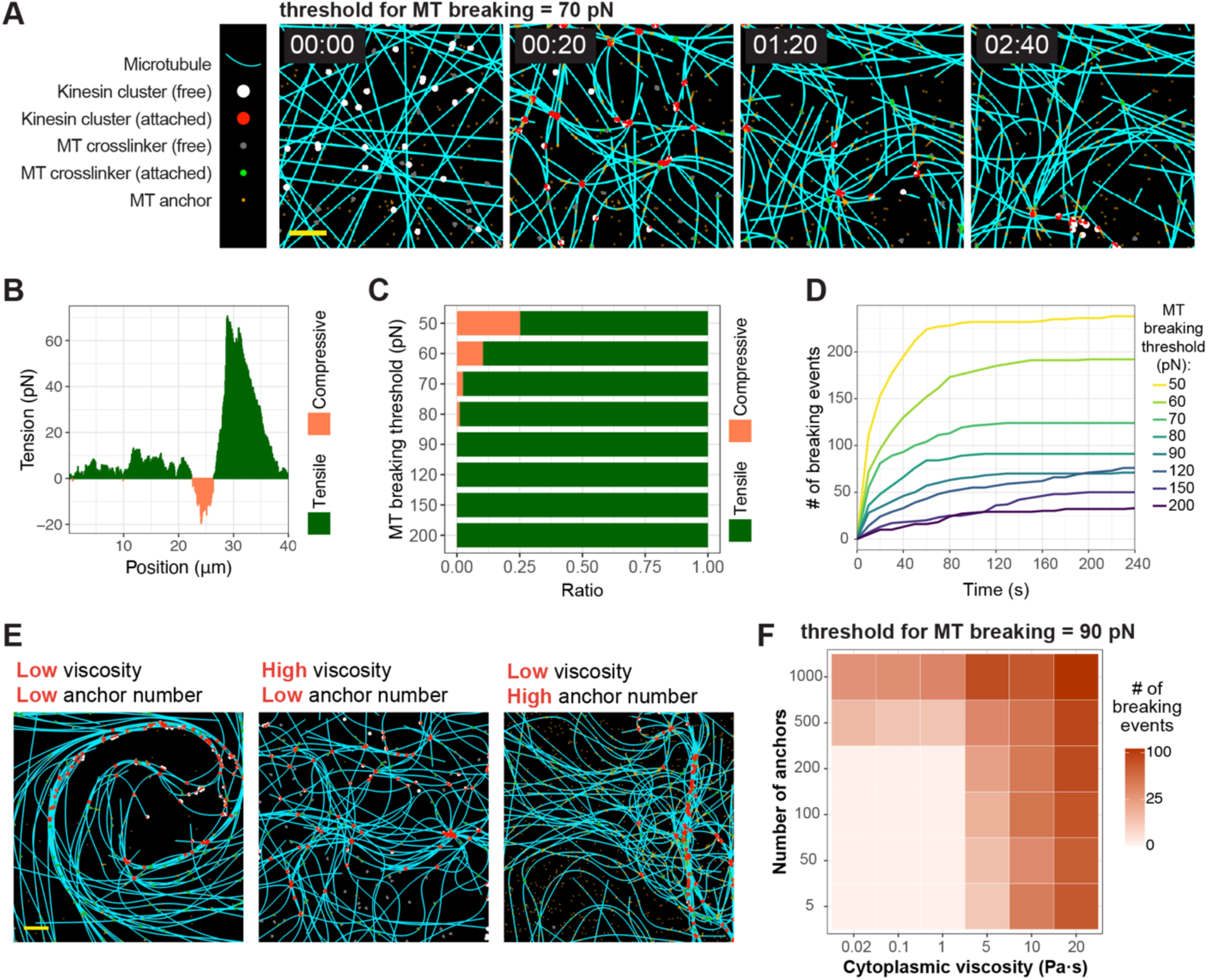
Cytosim simulation of the interactions between multi-kinesin clusters and the microtubule network. **(A)** An example Cytosim simulation run using the default values for all parameters and the threshold for microtubule breaking set to 70 pN. Cyan lines represent MTs. Circles represent multi-kinesin clusters that are not engaged (white color) or are engaged with microtubules (red color). Spots represent microtubule crosslinkers that are not engaged (gray color) or are engaged (green color) with microtubules. Orange dots represent microtubule anchors. Time stamp is min:sec. Scale bar (yellow): 2 μm. **(B)** Tension along the longitudinal axis of a representative microtubule in the simulation run shown in (A). X-axis is position along microtubule fiber; y-axis is the tension at each position. Tensile force is positive and colored green whereas compressive force is negative and colored orange. **(C)** The ratio of microtubule breaking events due to tensile or compressive forces at different threshold values of microtubule breaking. Microtubule breaking by tensile force is colored green, whereas breaking by compressive force is colored orange. Data are from one simulation run at each microtubule breaking threshold. **(D)** Number of microtubule breaking events over time at different threshold values for microtubule breaking. **(E)** Representative images of microtubule reorganization by multi-motor clusters at different values of cytoplasmic viscosity (low = 1 Pa·s, high = 10 Pa·s) and number of microtubule anchors (low = 50 anchors/400 μm^2^, high = 1000 anchors/400 μm^2^). Scale bar: 2 μm. **(F)** Heatmap showing the effect of changing cytoplasmic viscosity (x-axis) and number of MT anchors (y-axis) on microtubule breaking behavior with the threshold for microtubule breaking set to 90 pN.

**Table 1:** Parameters tested in Cytosim simulations.

| Parameter | Unit | Default value | Source | Tested range |
| --- | --- | --- | --- | --- |
| Motor processivity | $\mu\text{m}$ | 7 | This study | 0.7, 1.4, 2.3, 7, 70 |
| Motor stall force $F_{\text{st}}$ | pN | 4 | (Siddiqui et al., 2022) | 1, 4, 7, 10 |
| Motor unbinding force $F_{\text{ub}}$ | pN | 4 | | Set at $F_{\text{st}}$ |
| Motor velocity | $\mu\text{m} \cdot \text{s}^{-1}$ | 0.7 | This study | 0.01, 0.1, 0.5, 0.7, 1, 2 |
| Motor cluster stiffness | $\text{pN} \cdot \mu\text{m}^{-1}$ | 100 | | 5, 10, 20, 50, 100, 500 |
| Motor cluster number ($N_c$) | | 120 | | 2, 8, 30, 120 |
| Number of motors per cluster ($N_k$) | | 24 | | 2, 5, 10, 24, 60, 120 |
| Microtubule anchor number |  | 500 |  | 5, 50, 100, 200, 500, 1000 |
| Microtubule anchor unbinding rate | $\text{s}^{-1}$ | 0.01 | | 1, 0.1, 0.05, 0.01, 0.005, 0.001 |
| Cytoplasmic viscosity | $\text{Pa} \cdot \text{s}$ | 10 | (Yamada et al., 2000) | 0.02, 0.1, 1, 5, 10, 20 |
| Microtubule rigidity | $\text{pN} \cdot \mu\text{m}^2$ | 22 | (Gittes et al., 1993) | |
| Microtubule number |  | 60 |  |  |
| Microtubule crosslinker number |  | 100 |  |  |

An example of a simulation run using the default values for all parameters and with the threshold for microtubule breaking set to 70 pN is shown in Fig. 5 A. Over time, multi-motor clusters engage with microtubules (change from white to red) and cause microtubule reorganization due to the sliding of adjacent microtubules and their entanglement into high densities followed by microtubule breaking and fragmentation. During the run, an individual microtubule fiber experiences a mixture of tensile (stretching) and compression forces at different positions along its length (Fig. 5 B). Higher tension is generated when microtubules are under tensile force (Fig. 5B), presumably because microtubules under compression can dissipate force by bending. Indeed, most microtubule fibers break when they are subjected to tensile forces (Fig. 5 C).

We were particularly interested in understanding how much force a microtubule can withstand before breaking, an important microtubule parameter that has not been directly measured. We thus tested a range of threshold values and found that microtubules broke quickly when the threshold for breaking was set at 50 pN whereas few microtubule breaking events were observed when the threshold for breaking was set at 200 pN (Fig. 5 D). From all of the conditions tested in the simulations, a microtubule breaking threshold in the range of 70-120 pN appears to recapitulate microtubule network reorganization and microtubule fragmentation that are similar to our experimental observations (Fig. 4). This finding implies that although the force produced by individual KIF1C motors (4 pN) is well below the threshold for microtubule breaking, motors clustered into KIF1C condensates may be able collectively impose a force equal to or higher than the range of 70-120 pN.

### Impact of cytoplasmic viscosity and microtubule anchoring on breaking

A key parameter that profoundly affects microtubule breaking behavior in the simulations is cytoplasmic viscosity. It has been challenging to experimentally determine the viscosity of cultured cells (Luby-Phelps et al., 1986; Yamada et al., 2000; Kalwarczyk et al., 2011; Delarue et al., 2018), with results ranging from 0.004 Pa·s to ∼10 Pa·s (*i*.*e*. from 4 to 10,000 times more viscous than water). Even within one cell, the viscosity can be spatially heterogeneous as the lamellar region at the periphery of the COS-7 cells used in our experiments was shown to be more viscous than the perinuclear region (Yamada et al., 2000). In the simulations, at low viscosity (0.02 - 1 Pa·s), the microtubule network became reorganized into loops of heavily bundled microtubules and very few microtubule breaking events occurred (Fig. 5 E,F). This behavior is reminiscent of previous *in vitro* reconstitution studies of multimeric kinesin clusters that organize microtubules into asters or nematic networks, a process important for the organization of mitotic microtubules in dividing cells (Nedelec et al., 1997; Surrey et al., 2001; Roostalu et al., 2018). However, at higher viscosity, the microtubules became entangled by multi-motor clusters and numerous microtubule breaking events were observed (Fig. 5 E,F), which more closely matches what we observed in our experiments with multi-KIF1C clusters (Figs. 1-4).

We reasoned that higher viscosity provides a drag force that resists microtubule reorganization by motor clusters. That is, at low viscosity, mechanical stress does not accumulate as the microtubules are easily deformed whereas at high viscosity, mechanical stress cannot be dissipated through microtubule bending and deformation. As an alternative mechanism to alter the drag force in the system, we tested how the number of microtubule anchors impacts microtubule breaking. Indeed, we found that adding more microtubule anchors decreased the formation of large, bundled microtubule loops, creating a microtubule network that looks similar to experimental observations (Fig. 5 E), and resulted in an increase in microtubule breaking events even at low viscosity (Fig. 5 F). Together, we conclude that drag forces provided by a viscous cytoplasm and/or microtubule anchoring are key to the accumulation of mechanical stress on microtubules that leads to microtubule breaking and fragmentation. We speculate that intracellular structures such as the actin meshwork may provide such drag force to microtubules and contribute to motor-cluster-induced microtubule fragmentation.

### The motility properties of the kinesin motor domain impact microtubule breaking

Our experimental results showed that the KIF1C motor domain is required for multi-KIF1C clusters to break and fragment microtubules in cells (Fig. 2). To explore how the motility properties of the kinesin in the multi-motor clusters impact the generation of mechanical tension that leads to microtubule breaking, we carried out simulations varying the parameters of motor velocity, processivity, and stall force at different thresholds of microtubule breaking. The results of these simulations predict that a motor with faster speed, higher processivity, or larger stall force will produce more microtubule breakage events upon clustering (Fig. 6 A).

**Figure 6.**
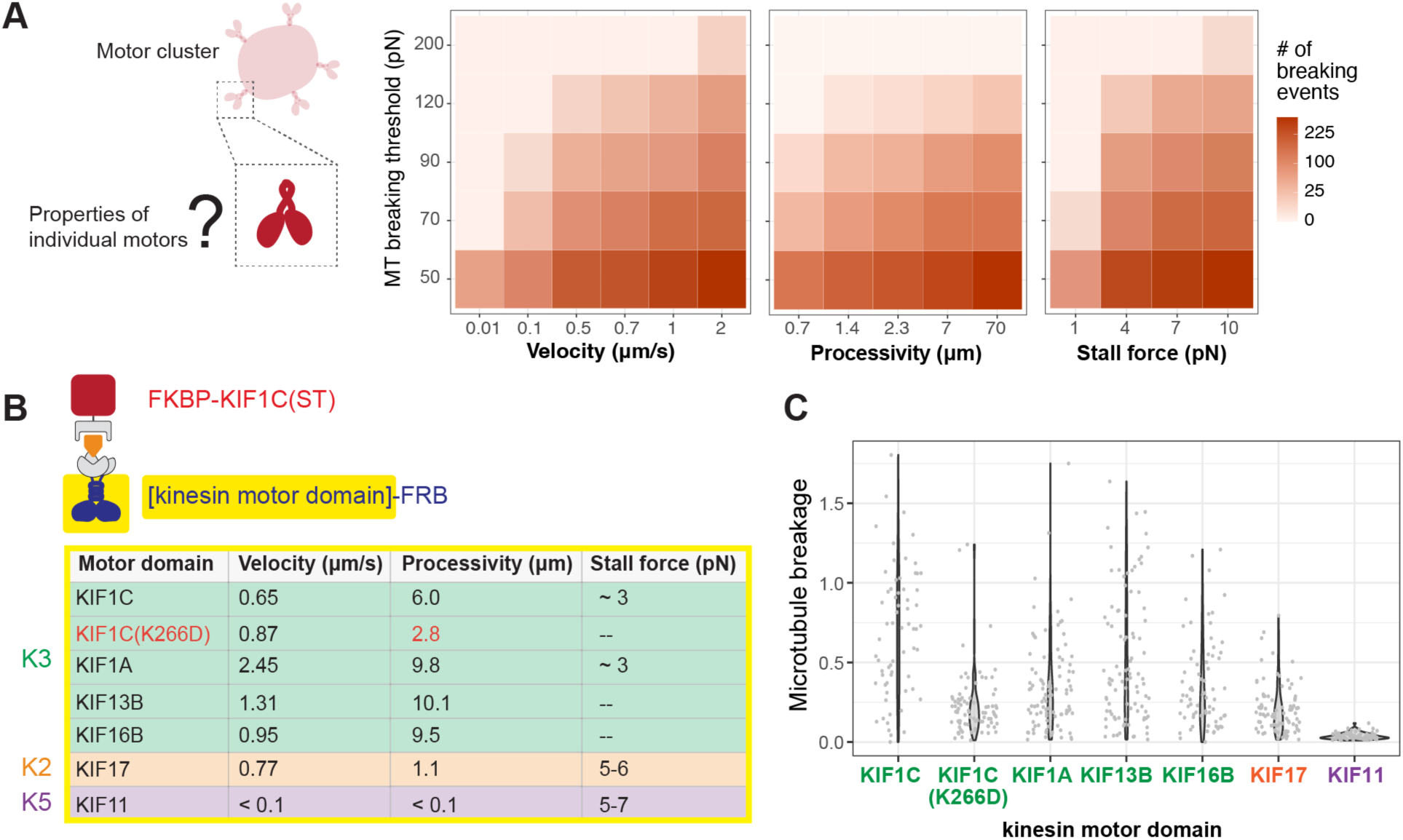
Impact of motor properties on microtubule breaking. **(A)** Heatmaps showing the effects of varying the biophysical properties of the kinesin motor domain (x-axes) in the cluster on microtubule breaking. Y-axis is the threshold force for microtubule breaking. **(B)** Schematic of the split-kinesin assay testing the impact of the biophysical properties of the kinesin motor domain on microtubule breaking. K3, K2, and K5 indicate kinesins belonging to the kinesin-3, kinesin-2, and kinesin-5 families, respectively. Values taken from literature (Valentine et al., 2006; Hammond et al., 2010; Soppina et al., 2014; Milic et al., 2017; Siddiqui et al., 2022) or this study (Fig. S4 A,B). **(C)** Violin plots showing quantification of microtubule breaking behavior in the split-kinesin assay with various kinesin motor domains and FKBP-KIF1C(ST) as the clustering mechanism. Each dot represents one cell. N = ∼80 cells per construct across 3 independent experiments.

To test these predictions experimentally, we performed the split-kinesin assay (Fig. 2 B) using KIF1C(ST) to mediate clustering of different kinesin motor domains in COS-7 cells (Fig. 6 B). We tested the motor domains of other kinesin-3s with higher velocities than KIF1C but similarly high processivities and low stall force, a kinesin-2 (KIF17) with similar velocity as KIF1C but lower processivity and higher stall force, and a kinesin-5 (KIF11) with lower velocity and processivity than KIF1C but higher stall force (Fig. 6 B). After 30 min of rapamycin treatment to induce the formation of multi-kinesin clusters, the cells were fixed and stained for β-tubulin to observe the microtubule network (Fig. S4). Quantification of microtubule breakage events shows that all of the kinesn-3 motor domains generate comparable levels of microtubule breaking when clustered into condensates despite their varying velocities (Fig. 6 C). In contrast, the KIF17 motor domain caused less microtubule breaking and the KIF11 motor domain resulted in little to no microtubule breaking when clustered into condensates (Fig. 6 C) despite their higher stall forces, in contrast to predictions of the simulations. Rather, the inability of KIF17 and KIF11 clusters to generate mechanical tension and cause microtubule breaking appears to be due to their lower processivity.

We speculate that higher motor processivity promotes microtubule breaking because it increases the time of engagement with a microtubule for each motor domain in the cluster, thereby increasing the efficiency of collective force production for the cluster. To further probe the contribution of motor processivity to the collective force output of motor clusters, we generated a variant of KIF1C(MD) with a mutation of lysine 266 to aspartate (K266D). For KIF1A(MD), this mutation significantly decreased processivity (∼6-fold) with minimal effect on velocity (Scarabelli et al., 2015). We used an *in vitro* single molecule motility assay to verify that the K266D mutation causes a similar decrease in processivity in KIF1C (Fig. S4 A,B). When utilized in the split-kinesin assay, the K266D mutant of KIF1C(MD) showed less microtubule reorganization and breakage than the WT motor domain (Fig. 6 C, Fig. S4 C). These results provide support for our hypothesis that motor processivity is a key factor in determining the collective force output that multi-motor clusters can impart to microtubules.

### The biophysical properties of the clustering domain impact microtubule breaking

Our experimental results also showed that clustering of KIF1C motors into multi-motor condensates is required for KIF1C to break and fragment microtubules in cells (Fig. 2). To explore the molecular properties required for multi-clusters to generate sufficient mechanical tension that leads to microtubule breaking, we carried out simulations that varied the stiffness of the clusters and the number of motors per cluster at different thresholds of microtubule breaking. The results predict that a stiffer (less fluid) clustering mechanism leads to more microtubule breakage events (Fig. 7 A, left graph). Surprisingly, even a low level of stiffness (5 pN/μm) leads to some microtubule breakage, suggesting that even the liquid condensates of KIF1C have sufficient stiffness to generate tension in the system. The simulation results also predict that a higher copy number of motors on each cluster will produce more microtubule breakage events (Fig. 7 A, right graph).

**Figure 7.**
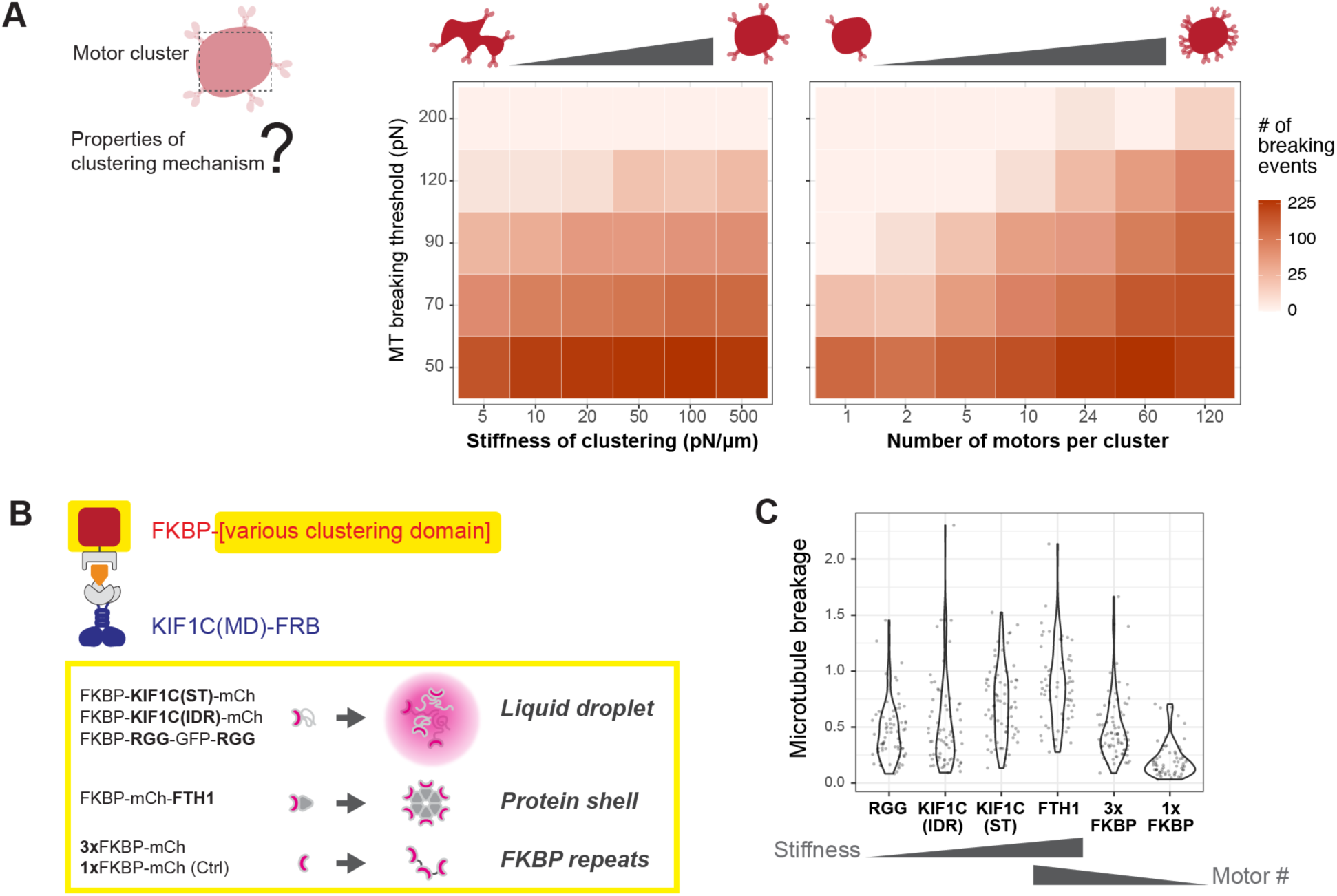
Impact of cluster properties on microtubule breaking. **(A)** Heatmaps showing the effects of varying the cluster properties on microtubule breaking. X-axes are the values tested for cluster stiffness (left graph) and number of motors (right graph); y axis is the threshold force for microtubule breaking. **(B)** Schematic of the split-kinesin assay testing the impact of the clustering mechanism on microtubule breaking. For clustering stiffness, domains forming condensates of various liquidity were compared to the FTH1 protein shell and for motor number per cluster, single and three tandem FKBPs were compared to the 24-mer FTH1 shell. **(C)** Violin plots showing quantification of microtubule breaking behavior in the split-kinesin assay with various clustering domains and the KIF1C(MD). Each dot represents one cell. N = ∼40 cells per construct across 3 independent experiments.

To test these predictions experimentally, we used the split-kinesin assay (Fig. 2 B) and varied the molecular mechanism used to generate clusters of the KIF1C(MD) (Fig. 7 B). To test the contribution of cluster stiffness, we compared the effects of clustering via three phase-separating condensates of varying liquidity to clustering via the solid-like FTH1 protein shell (Fig. 7 B). For the phase-separating condensates, we utilized the KIF1C(ST) (Fig. 2 A), the KIF1C(IDR) (Fig. 1 C; Geng et al., 2024) and a RGG repeat domain from the *C. elegans* protein LAF-1 that forms condensates in mammalian cell lines (Elbaum-Garfinkle et al., 2015; Schuster et al., 2018). The KIF1C(IDR) and RGG condensates are liquid-like as they show rapid fluorescence recovery kinetics in FRAP (fluorescence recovery after photobleaching) experiments [t_1/2_ recovery ∼ 1 sec (Schuster et al., 2018; Geng et al., 2024)]. In contrast, the KIF1C(ST), which contains both coiled-coil and IDR regions, forms less liquid condensates as indicated by slower fluorescence recovery kinetics [t_1/2_ recovery ∼ 17 seconds (Geng et al., 2024)].

Clustering of the KIF1C(MD) via the IDR or FTH1 domains was induced by rapamycin treatment and the microtubule network was visualized by staining for β-tubulin. Quantification of microtubule breakage events shows that clustering via either of the liquid-like phase-separating condensates [KIF1C(IDR) or RGG] leads to less microtubule breakage than clustering via the less liquid KIF1C(ST) or the solid FTH1 shell (Fig. 7 C, Fig. S5, Video 3). Indeed, the more liquid-like nature of KIF1C(IDR) and RGG can be observed in split-kinesin assay, as their spherical droplet shape is transformed to filament-like shape decorating microtubules upon rapamycin treatment, whereas KIF1C(ST) condensates remain spherical after rapamycin treatment (Fig. S5, Video 3). These results are consistent with the simulation results (Fig. 7 A) which predicted that clustering via more liquid-like condensates (lower cluster stiffness) results in fewer microtubule breaking events.

To test the contribution of motor number per cluster, we compared the effects of clustering via the FTH1 protein shell to clustering via single or tandem FKBP domains. The self-assembling FTH1 protein shell provides binding sites for up to 24 KIF1C(MD)s whereas three tandem FKBP domains (3xFKBP) can generate a cluster of three KIF1C(MD)s and a single FKBP domain serves as a control as it allows no clustering of KIF1C(MD)s. Clustering of even small numbers of KIF1C(MD) by the 3xFKBP construct resulted in localized microtubule bundling and breakage (Fig. S5). Quantification of microtubule breakage shows that clusters with higher numbers of motors (i.e., FTH1) result in more microtubule breakage than clusters with lower numbers of motors (i.e., 3xFKBP) (Fig. 7 C). These results are consistent with the simulation results (Fig. 7 A), which predicted that motor copy number per cluster is an important contributor to the effects of multi-motor clusters on microtubule organization and integrity.

### The kinesin-cargo interface of KIF16B can protect microtubules from motor-cluster-induced mechanical stress

Our experimental data and computational modeling results demonstrate that microtubules are susceptible to mechanical stress imposed by multi-kinesin clusters. This is surprising as multi-kinesin clusters exist in biologically-relevant contexts yet the microtubule integrity does not appear to be compromised. Indeed, there are many examples in the literature where multiple kinesin motors are present on the surface of membrane-bound organelles, presumably to enable efficient transport within the complex cellular environment (Jiang et al., 2019; Sarpangala and Gopinathan, 2022; Bensel et al., 2024; Jiang et al., 2024). It is therefore important to ask why motors clustered on the surface of vesicles and organelles do not induce sufficient mechanical stress on microtubules that leads to their breaking and fragmentation.

As an example, the motility of early endosomes is driven by multiple kinesin-1 and kinesin-3 motors (Nagpal et al., 2024), yet their motility does not alter the microtubule network or cause any microtubule breakage (Video 4). Even overexpression of an early endosome transporter, the kinesin-3 KIF16B, did not cause reorganization or breakage of the surrounding microtubules (Fig. 1 D). KIF16B associates with early endosomes via binding of its C-terminal PX domain to PI(3)P (phosphatidylinositol-3-phosphate) lipid molecules in the membrane (Fig. 8 A) (Hoepfner et al., 2005; Blatner et al., 2007). We speculated that KIF16B’s protein-lipid mechanism of cargo attachment may be a critical feature that prevents the buildup of mechanical stress in the cargo-motor-microtubule system. To test this possibility, we generated a chimeric construct [1C(MD)-16B(PX), Fig. 8 B] in which the KIF1C(MD) (aa 1-376) is fused to the KIF16B PX domain (aa 1179-1317). The chimeric construct showed strong localization to endosomal cargoes when expressed in COS-7 cells, however, the microtubules remained intact despite the reorganization of the surrounding microtubule network into a higher density by the KIF1C motors clustered on the surface of early endosomes (Fig. 8 B). These results indicate that multi-motor clusters in which kinesin-3 motor domains (KIF16B in the native molecule or KIF1C in the chimeric molecule) are attached to a membrane surface via a protein-lipid interaction do not induce microtubule tension that leads to breakage and fragmentation.

**Figure 8.**
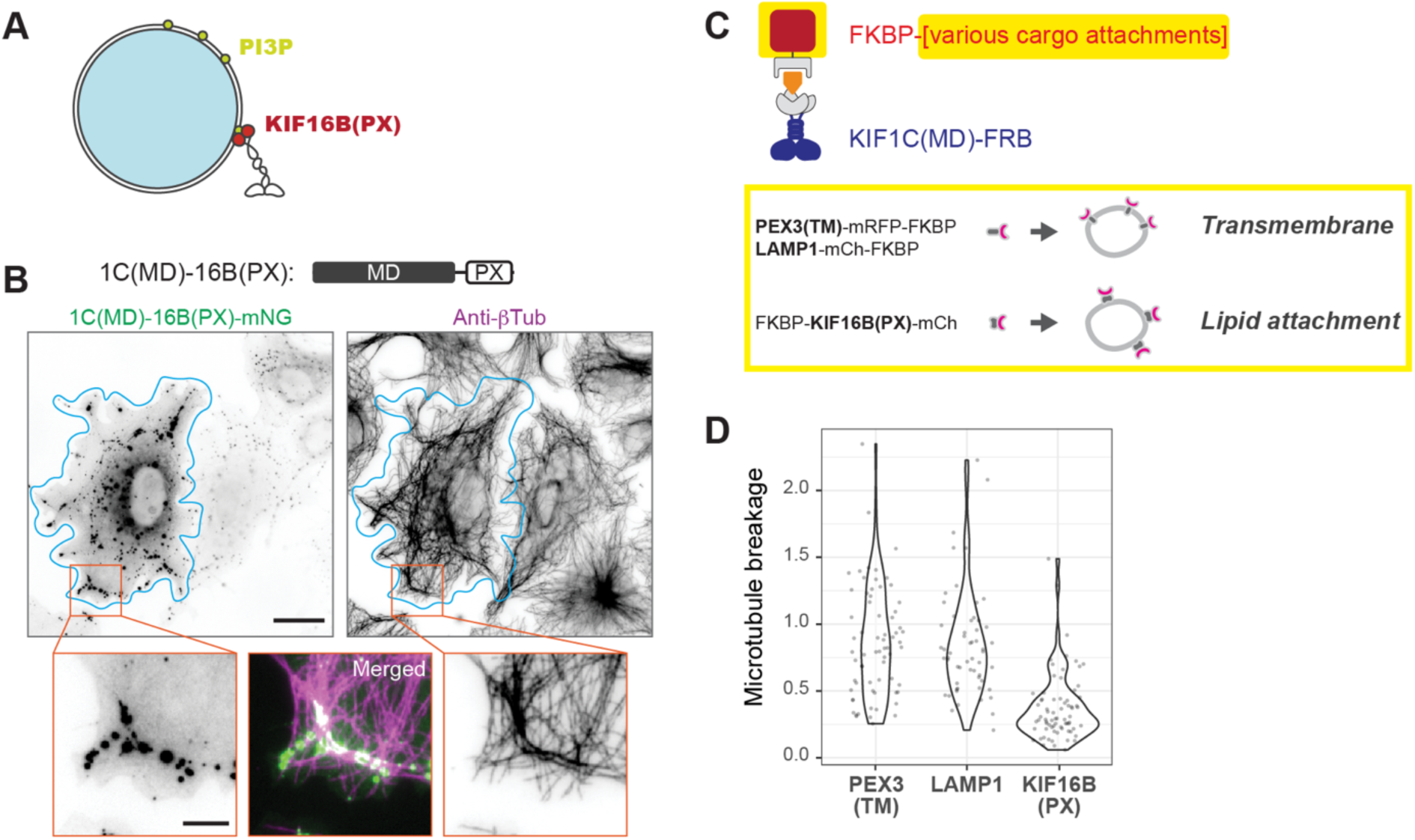
KIF16B PX domain mediates multi-kinesin clustering but prevents microtubule breakage. **(A)** Schematic showing attachment of the kinesin-3 KIF16B to endosomal membranes via the interaction of its PX domain with PI(3)P lipid molecules. **(B)** Schematic of the 1C(MD)-16B(PX) chimeric protein containing the KIF1C(MD) fused to the KIF16B PX domain. Representative immunofluorescence images of COS-7 cells expressing 1C(MD)-16B(PX) and fixed and stained for microtubules (anti-βTub). Cyan lines mark the cell outline and orange boxes indicate the regions shown in the magnified images. Scale bars: 20 μm for whole-cell views and 5 μm for magnified images. **(C)** Schematic of the split-kinesin assay testing the impact of motor-cargo attachment mechanism on microtubule breaking. The transmembrane domains of PEX3 or LAMP1 or the protein-lipid attachment of KIF16B(PX) were used to generate clusters of KIF1C(MD). **(D)** Violin plots showing quantification of microtubule breaking behavior in the split-kinesin assay with motor-cargo attachment mediated by the transmembrane (TM) domains of PEX3 or LAMP1 or the KIF16B(PX) domain. N = ∼40 cells per construct across 3 independent experiments.

Why is motor clustering mediated by KIF16B(PX) unable to generate sufficient tension in the system to cause microtubule breaking? A potential clue comes from a previous study that measured the attachment of KIF16B(PX) to PI(3)P-containing liposomes using optical tweezers. The lifetime of the bonds between KIF16B(PX) and PI(3)P decreased by ∼ 20-fold (from 390 ms to 20 ms) in response to increased load (from 11 pN to 45 pN), indicating slip-bond behavior (Pyrpassopoulos et al., 2017). We hypothesized that a protein-lipid interaction for motor-cargo attachment prevents mechanical stress from building up in the cargo-motor-microtubule system since KIF16B(PX) will dissociate from the cargo membrane at lower forces (11-45 pN) than the threshold force for microtubule breaking (70-120 pN).

To test this hypothesis, we used the split-kinesin assay to vary the motor-cargo attachment mechanism. We investigated the impact on microtubule breaking for KIF1C(MD) organized into multi-kinesin clusters via the KIF16B(PX) as compared to the transmembrane (TM) domains of LAMP1 (lysosomal protein) and PEX3 (peroxisomal protein) (Fig. 8 C). Transmembrane proteins are stably integrated into a lipid bilayer and require forces as high as 200-500 pN to dissociate from the membrane (Fotiadis, 2012; Petrosyan et al., 2015), which is higher than the threshold force for microtubule breaking. Rapamycin treatment induced the formation of multi-KIF1C clusters on early endosomes [via KIF16B(PX)], lysosomes (via LAMP1), or peroxisomes [via PEX3(TM)] and the cells were fixed and stained for β-tubulin to visualize the microtubule network. Formation of multi-KIF1C clusters via transmembrane segments strongly induced microtubule fragmentation (Fig. S6). Quantification of these results demonstrates that multi-motor clusters formed by transmembrane segments induced more microtubule fragmentation that those formed by the lipid-associated KIF16B(PX) domain (Fig. 8 D). These results support our hypothesis that the labile motor-cargo interface of KIF16B can prevent the accumulation of mechanical stress in the cargo-motor-microtubule system that leads to microtubule breakage.

## Discussion

We show that self-association of the kinesin-3 motor KIF1C into biomolecular condensates results in multi-motor clusters that entangle neighboring microtubules and cause their breakage and fragmentation in cells. Although an artifact of KIF1C overexpression, this system enabled us to study how microtubules respond to mechanical stress within the cargo-motor-microtubule system (Fig. 9). Using a combination of immunofluorescence, live-cell imaging, and computational modeling, we show that motor processivity, clustering mechanism, and drag force are key parameters that impact the system response. We estimate a microtubule rupture force of 70-120 pN in cells, which is lower than previous estimates based on *in vitro* studies with taxol-stabilized microtubules.

**Figure 9.**
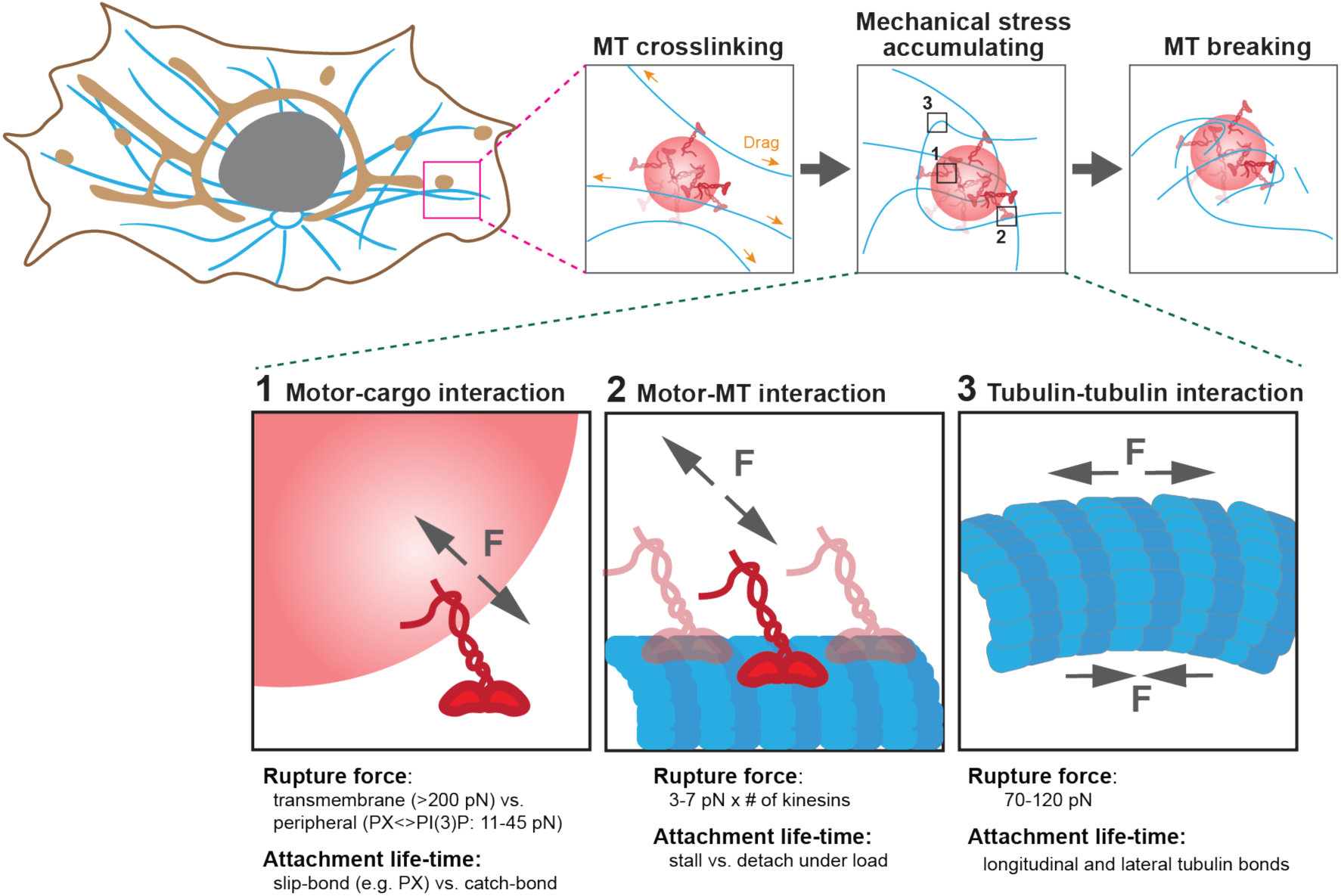
Model for how the buildup of mechanical stress in the cargo-motor-microtubule system can be relieved. Individual kinesin motors within a multi-motor cluster are able to walk along and crosslink multiple microtubule tracks. Given the high drag force imposed within the cytoplasm, mechanical stress accumulates in the system. The relative strengths of the three types of interactions in the cargo-motor-microtubules system (bottom) determines whether stress is relieved by microtubule breaking. In most cases, the the kinesin-cargo interaction (box 2) or the kinesin-microtubule interaction (box 2) will release before rupture of the tubulin-tubulin interactions within the microtubule (box 3). For each attachment point, the interactions are governed by the amount of force that they can withstand and the lifetime of interaction under force.

### Multi-motor clusters generate mechanical stress that leads to microtubule breakage

Our observations reveal that multi-motor clusters can generate mechanical stress that leads to microtubule breakage and fragmentation. We suggest that motors in the cluster can generate collective forces that are high enough to rupture tubulin-tubulin interactions. An alternative possibility is that high microtubule curvature induced by multi-kinesin clusters promotes microtubule breakage indirectly by recruiting cytoplasmic severing enzymes, such as katanin and spastin (Odde et al., 1999; McNally and Roll-Mecak, 2018). We do not currently favor this model as conditions that generate highly curved microtubules in cells, such as treatment with the kinesin-1 activator kinesore (Randall et al., 2017) or expression of superprocessive kinesin-3 motors (Budaitis et al., 2022), do not result in microtubule breakage and fragmentation. In addition, tightly buckled microtubules break in the absence of severing factors in *in vitro* experiments (Kabir et al., 2020; Tsitkov et al., 2022; Nasrin et al., 2024).

From the simulations, we find that most microtubule fibers are broken by tensile forces rather than compression forces (Fig. 5 B and C), consistent with previous *in vitro* studies (Kabir et al., 2014, 2015, 2020; Nasrin et al., 2024). Although longitudinal interactions between tubulin subunits are stronger than the lateral interactions between protofilaments, microtubules are more susceptible to tensile forces since microtubules under compression can dissipate force by bending.

We also find that more microtubule breakage is observed under high viscosity conditions or in the presence of anchors to the substrate (Fig. 5 E and F). We surmise that including microtubule anchors or a high viscosity help motor clusters to exert force on microtubules by providing drag. This is consistent with previous work where microtubule pinning leads to an increase in microtubule breaking (Kabir et al., 2020; Tsitkov et al., 2022).

Using the split-kinesin assay, we show that both the KIF1C motor domain and a clustering mechanism are required for the generation of mechanical stress that leads to microtubule breakage. For the motor domain, processivity is a key property as kinesin motor domains with high processivity generate more breakage and a mutation that decreases KIF1C processivity results in less breakage (Fig. 6 and S4). We hypothesize that processivity is critical as it defines how long an individual motor domain maintains an interaction with the microtubule, and thus determines the time window in which multiple motors in a cluster can cooperate to impact microtubule structure and mechanics. Consistent with this, motor processivity was found to be a key variable determining the size of the microtubule aster formed by dimeric and tetrameric motors (Banks et al., 2023).

We were initially surprised that the IDR-driven formation of liquid condensates would generate sufficient mechanical stress that leads to microtubule rupture. Indeed, KIF1C condensates are dynamic structures that undergo fission and fusion (Geng et al., 2024). We surmise that the large number of active motors in the condensate likely compensates for the tendency of the condensate to dissolve, as motor number per cluster was found to be a critical factor in the simulations. For the clustering mechanism, we find that a stiffer clustering mechanism can impart greater mechanical stress as a denser condensate (i.e., the KIF1C ST) (Geng et al., 2024) or a protein shell (i.e., FTH1 24-mer) leads to more breakage events.

### Limitations and caveats to the computational modeling of microtubule breaking

Computational modeling using Cytosim provided a mechanistic understanding of microtubule breaking in response to entanglement by multi-kinesin clusters, however, there are several limitations to our approach. First, our simulations primarily considered tensile and compressive stresses as a determinant of microtubule breaking behavior, whereas other types of mechanical stress such as shear force and torsion could not be evaluated. We focused on tensile stress based on previous atomistic molecular dynamics simulations suggesting that microtubule breaking is more sensitive to stretching over indentation or dilation (Wu and Adnan, 2018). However, torsion may have a significant impact on microtubule breaking. For actin filaments, it has been shown that the application of twisting before stretching decreased the tensile force needed to break a single actin filament (Tsuda et al., 1996). For microtubules, motor proteins can generate torque by helical motion along microtubules (Meißner et al., 2024) and the sensitivity of microtubules to such torsion will be important to address in future work.

Second, our simulations did not consider the possibility of material fatigue of microtubules under repetitive stress. Material fatigue is common in materials such as metals and plastics where a force smaller than the ultimate rupture force can cause fracturing if it is applied in a cyclic fashion (Salvati, 2024). Recent *in vitro* studies suggest that microtubules indeed display properties of material fatigue (Schaedel et al., 2015; Nasrin et al., 2024). It seems likely that the repeated deformation of microtubules around multi-motor clusters (Fig. 3 and S2; Video 1, 2 and 3) will lead to material fatigue and lower forces for microtubule rupture over time.

### Comparison to previous in vitro estimates of microtubule rupture force

Our experiments and simulations suggest that microtubules in cells are surprisingly fragile as they break when subjected to mechanical stress produced by clusters of kinesin motor proteins. We estimate a rupture force of 70-120 pN in cells, a fundamental microtubule feature that has not previously been measured. However, our rupture forces for microtubule in cells are lower than previous estimates from *in vitro* studies (Endow and Marszalek, 2019; Kabir et al., 2020). An important consideration is that the *in vitro* experiments necessarily used taxol to stabilize microtubules and prevent their depolymerization from the ends. However, taxol is also known to soften microtubules and allows them to withstand buckling at radii of curvature that correlate with microtubule breaking in cells (Kikumoto et al., 2006; Hawkins et al., 2013; Lopez and Valentine, 2014; Kabir et al., 2015, 2020; Tsitkov et al., 2022).

An additional consideration is that microtubules in cells can incur damage from molecular motors walking along the lattice surface, as well as from microtubule crossings or collisions (Aumeier et al., 2016; de Forges et al., 2016; Triclin et al., 2021; Andreu-Carbó et al., 2022; Budaitis et al., 2022). Damage to or defects in the microtubule lattice lead to increased material fatigue (Schaedel et al., 2015, 2019) and make them more susceptible to breaking (Jiang et al., 2017; Budaitis et al., 2022).

### Relevance for multi-motor transport in cells

Our finding that microtubules are surprisingly prone to breaking is especially striking when one considers the physiological role of microtubules in supporting intracellular transport events where the presence of multiple motor proteins on a cargo is common (Ashkin et al., 1990; Jiang et al., 2019; Sarpangala and Gopinathan, 2022; Bensel et al., 2024; Jiang et al., 2024). We suggest that cells have evolved mechanisms to protect microtubule integrity in the context of cargo-motor-microtubule systems (Fig. 9). We propose that either the motor-cargo interaction (Fig. 9, box 1) or the motor-microtubule interaction (Fig. 9, box 2) ruptures to relieve mechanical stress in the system before the microtubule breaks (Fig. 9, box 3).

For the motor-cargo interaction (Fig. 9, box 1), the tension-dependent detachment of a motor from the cargo surface has not been studied but is likely to be a key parameter is dissipating mechanical stress in the system. Our simulations and experiments have shown that increasing the strength of the motor-cargo interaction (via transmembrane domains) results in more microtubule breakage whereas decreasing the strength of the motor-cargo interaction [via the KIF16B(PX) domain] leads to less microtubule breakage. A key parameter is the lifetime of the bonds under force and in this respect, it is intriguing that the KIF16B(PX) domain behaves like a slip bond in that it gives up upon increase of tension (Pyrpassopoulos et al., 2017). The slip-bond design of kinesin-cargo interface might be a general mechanism to prevent tension building up on microtubules while allowing efficient cargo transport.

For the motor-microtubule interaction (Fig. 9, box 2), individual kinesin motors are known to detach from the microtubule when subjected to 3-7 pN of hindering force. It thus seems likely that individual motors within a cluster will detach from the microtubule before mechanical stress builds to the 70-120 pN of force needed to rupture a microtubule. However, some kinesins are known to stall under force (e.g., kinesin-1) whereas others rapidly detach under force (e.g., kinesin-3) and thus the lifetime of attachment to the microtubule under force is likely to be a key parameter that impact the dissipation of mechanical stress in the system.

In addition, there are other strategies that can be employed by cells to prevent multi-motor clusters from generating high mechanical stress in the cargo-motor-microtubule system. For KIF1C, the low concentration of endogenous KIF1C protein restricts condensate formation to small structures located at cell protrusions where there are limited microtubules for engagement (Geng et al., 2024). For other kinesins, autoinhibition is a well-known mechanism (Verhey and Hammond, 2009) that may be applicable to motors present on a cargo surface. Finally, proteins such as kinesin binding protein (KIFBP) regulate the interaction of kinesins with microtubules (Kevenaar et al., 2016) and could dampen the impact of multi-motors present on a cargo surface.

### Kinesin clustering and microtubule breaking as an intracellular force probe

In our split-kinesin assay, we found that a variety of intracellular materials (biomolecular condensates, organelle-associated proteins, self-assembled protein shell) can mediate the clustering of motor proteins and induce variable levels of microtubule fragmentation. Therefore, we propose that our split-kinesin system can be repurposed to probe the mechanical properties of different molecular assemblies. For example, one such application is to measure the liquidity of different phase-separated condensates. By measuring the level of microtubule fragmentation, one can qualitatively assess the mechanical strength of molecular assemblies in the cytoplasm of live cells in a non-invasive manner.

## Materials and methods

### Cell culture

COS-7 [male Ceropithecus aethiops cells (African green monkey) kidney fibroblast, RRID: CVCL_0224] were grown in in Dulbecco’s Modified Eagle Medium (Gibco) supplemented with 10% (vol/vol) Fetal Clone III (HyClone) and 2 mM GlutaMAX (L-alanyl-L-glutamine dipeptide in 0.85% NaCl, Gibco). hTERT-RPE1 cells (female Homo sapiens retinal pigment epithelium, RRID: CVCL_4388) were grown in DMEM/F12 with 10% (vol/vol) FBS (HyClone), 0.5 mg/ml hygromycin B, and 2 mM GlutaMAX (Gibco). Both cell lines were purchased from American Type Culture Collection and grown at 37°C with 5% (vol/vol) CO_2_. Both cell lines are checked annually for mycoplasma contamination and COS-7 cells were authenticated through mass spectrometry (the protein sequences exactly match those in the *Ceropithecus aethiops* genome).

### Plasmids and Adenoviral vectors

The KIF1C motor domain (MD) contains the first 376 amino acids of human KIF1C, a short segment of neck coil from KIF1A (aa sequence LYAQGLGDITDGAG), followed by a leucine zipper sequence (aa sequence VKQLEDKVEELASKNYHLENEVARLKKLV) to ensure dimerization. The motor domain constructs of other kinesins, including KIF1A(MD), KIF13B(MD), KIF16B(MD), KIF17(MD) and KIF11(MD) are from previous studies (Hammond et al., 2010; Soppina et al., 2014). Some of the plasmids were subcloned to change their fluorescent protein tags. Because the fluorescence intensity of KIF1C(MD)-tagBFP2-FRB was weak and difficult to quantify, a KIF1C(MD)-FRB-IRES-H2B-tagBFP2 construct was generated and utilized (Fig. S5 and S6) to drive the expression of two peptides, KIF1C(MD)-FRB and H2B-tagBFP2 under the same promoter. The concentration of the tagBFP2 signal in the nuclei through the localization of histone H2B enables easier quantification of fluorescence intensity, which was used as an indicator of KIF1C(MD)-FRB expression level in those cells.

PEX3(TM)-mRFP-FKBP is described previously (Budaitis et al., 2019). Other FKBP-tagged constructs were cloned by inserting DNA fragments (IDR, gBlocks Gene Fragments) encoding FKBP at the ends of various constructs to generate FKBP-KIF1C(ST)-mCherry and FKBP-KIF1C(IDR)-mCherry (Geng et al., 2024), FKBP-RGG-GFP-RGG (Addgene #124930) (Schuster et al., 2018), FKBP-KIF16B(PX)-mCherry (this study), LAMP-mCherry-FKBP (Qian et al., 2009), and FKBP-mCherry-FTH1 (Addgene #100749) (Clarke and Royle, 2018). mCherry-EEA1 is cloned from GFP-EEA1 (AddGene #42307) (Lawe et al., 2000).

Plasmids for expression in mammalian cells utilize the cytomegalovirus (CMV) promoter. The adenovirus plasmid encoding EGFP-*α<u>-</u>*tubulin (pShuttle-EGFP-tubulin) was a gift from Torsten Wittmann (Addgene plasmid #24327, RRID: Addgene_24327) (Kumar et al., 2009) and adenovirus was produced by the University of Michigan Vector Core. All plasmids were verified by whole-plasmid DNA sequencing (Plasmidsaurus).

### Transfection and immunofluorescence

For transfection of cells in 6-well plates or 35-mm dishes (2 mL final volume of media each well/dish), 1 μg plasmid DNA and 3 μL TransIT-LT1 transfection reagent (Mirus #2300) were diluted in 100 μL Opti-MEM Reduced-Serum Medium (Gibco #31985062) to make a transfection mixture which was added to media in each well immediately after seeding cells (0.5-1 × 10^5^ cells). For split kinesin assay where various clustering domain constructs were co-transfected with KIF1C motor domain (Fig. 7 and Fig. S5), 350 ng DNA was used for KIF1C motor domain constructs while 1 μg DNA was used for each clustering domain construct.

16-24 h post transfection, the cells were fixed and permeabilized in pre-chilled methanol for 8 min at −20 °C, followed by blocking with 0.2% FSG (fish skin gelatin) in PBS for 5 min. Primary antibodies (Ms anti-β-tubulin, DSHB #E7, 1:2,000 dilution) were applied in 0.2% FSG in PBS overnight at 4 °C in a custom humidified chamber. After 3 washes with 0.2% FSG in PBS, secondary antibodies were applied in 0.2% FSG in PBS for 1 h at room temperature in the dark. Nuclei were stained with 10.9 mM 40,6-diamidino-2-phenylindole (DAPI) and the coverslips were mounted using Prolong Gold (Invitrogen #P36930). Images were acquired on an inverted epifluorescence microscope (Nikon TE2000E) with a 60×, 1.40 NA oil-immersion objective and a CoolSnap HQ camera (Photometrics).

### Imaging of live cells

Cells were seeded in 35 mm glass-bottom dishes (Matek #P35G-1.5-14-C) and were transfected as described above. 16–24 h post-transfection, cells were infected with EGFP-α<u>-t</u>ubulin adenovirus for another 16-24 h. The cells were washed with and then incubated in Leibovitz’s L-15 medium (Gibco #11415064) and imaged at 37 °C in a temperature-controlled and humidified stage-top chamber (Tokai Hit) on a Nikon X1 Yokogawa Spinning Disk Confocal microscope with a 60x, 1.49 NA oil-immersion objective, and a Andor DU-888 camera. 500x rapamycin stock in ethanol was added into the media to reach a final concentration of 44 nM. Video recording started 10-15 min after the addition of rapamycin and lasted for 4 min with an acquisition speed of 2 frames per second. Image acquisition was controlled with Elements software (Nikon). Images were processed in Fiji to generate videos and time series images. For visual clarity, zoomed-in images of subcellular regions in Fig. 3 and Fig. S2 were smoothened by the “Scale” function of Fiji to scale up by a factor of 4.

### Cell lysate preparation

24 h after transfection, COS-7 cells were harvested and pelleted by centrifugation at 3000 x g at 4°C. The cell pellet was washed with PBS and resuspended in ice-cold lysis buffer (25 mM HEPES/KOH, pH 7.4, 115 mM potassium acetate, 5 mM sodium acetate, 5 mM MgCl_2_, 0.5 mM EGTA, and 1% [vol/vol] Triton X-100) freshly supplemented with 1 mM ATP, 1 mM PMSF, and protease inhibitor cocktail (Sigma-Aldrich #P8340). After clearing the lysate by centrifugation for 10 min at 20,000 x g at 4°C, aliquots were snap frozen in liquid nitrogen and stored at −80°C until further use.

### Single molecule motility assays with TIRF microscopy

A flow cell (∼10 μl volume) was assembled by attaching a clean #1.5 coverslip to a glass slide with two thin strips of parafilm. The glass slide was briefly placed on a hotplate to slightly melt the parafilm strips, creating a seal between the coverslip and the slide. Taxol-stabilized microtubules were assembled as previously described (Norris et al, 2015). HiLyte-647-labeled microtubules were polymerized from purified tubulin (Cytoskeleton #TL670M) in BRB80 buffer (80 mM PIPES/KOH, 1 mM EGTA, and 1 mM MgCl_2_, pH 6.8) supplemented with 1 mM GTP at 37°C for 15 min. After 1:6 dilution by prewarmed BRB80 containing 20 μM taxol and an additional 35 min incubation at 37°C, polymerized microtubules were stored at room temperature in the dark for up to a week. Before imaging, polymerized microtubules were further diluted in BRB80 buffer containing 20 μM taxol, then infused into a flow cell and incubated for 5 min at room temperature to attach to the coverslip surface. Subsequently, blocking buffer (10 mg/mL bovine serum albumin (BSA) in BRB80 buffer with 10 μM taxol) was infused into the flow-cell and incubated for 3 min to prevent nonspecific binding to the coverslip surface. Finally, lysates containing equal amounts of motor proteins (typically 0.1–1.0 μL) were added to the flow chambers in a motility mixture in BRB80 buffer. Each motility mixture also contained 2 mM ATP, 10 mg/mL BSA, 10 μM taxol, and oxygen-scavenging components (1 mM dithiothreitol, 1 mM MgCl_2_, 10 mM glucose, 0.1 mg/mL glucose oxidase, and 0.08 mg/mL catalase) to reduce photobleaching.

The flow cells were sealed with molten paraffin wax. Imaging was performed on an inverted Ti-E/B TIRF (Total Internal Reflection Fluorescence) microscope equipped with a perfect focus system, a 100x, 1.49 NA oil immersion TIRF objective, three 20-mW diode lasers (488 nm, 561 nm, and 640 nm) and an electron-multiplying charge-coupled device detector. Image acquisition was controlled with NIS-Elements software (Nikon). Multi-channel time-lapse images were acquired at a rate of every 500 ms for 5 min. The temperature of the microscope room was set at 23°C. Images and videos were processed in Fiji (https://fiji.sc/). Kymographs of single-molecule assay for kinesin motility were generated using the Multi Kymograph function in Fiji and were used to measure the velocity and run length of kinesins.

### Simulation of microtubule breaking by Cytosim

The simulation of microtubule breaking by clustered motors was done utilizing Cytosim, an open-source cytoskeleton simulation project developed by the Nédélec group (https://gitlab.com/f-nedelec/cytosim). In the simulation, microtubules in a 2-dimentional space are represented by fiber objects that are composed of small segments connected by vertices. Cytosim models the microtubules as mechanical objects in a low Reynolds number regime where inertia is negligible. The motion of such objects and the forces they experience are described by Langevin dynamics (Nedelec & Foethke, 2007). To enable the breaking of microtubule fibers, the source code was modified with help from F. Nédélec for comparison of the force applied to each segment of the fiber as compared to a user-defined threshold of breaking force. The breaking behavior of microtubules is controlled by the following equation:

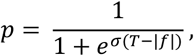

where *p* is the probability of breaking; *T* is the threshold force for microtubule breaking defined by the user in the configuration files; *f* is the highest force across all segments of the current microtubule fiber (a positive *f* value indicates tensile (stretching) force, a negative *f* value indicates compressive force); *σ* controls how sensitive the change of probability *p* is to the change of forces near the threshold *T*.

To test motor processivity, the parameter processivity (μm) in the simulation was modified indirectly by changing the motor unbinding rate (s^-1^) parameter by a range of (1, 0.5, 0.3, 0.1, 0.01), while keeping the motor velocity parameter constant (0.7 μm·s^-1^). Since motor processivity = velocity/unbinding rate, the range of processivity parameter tested is (0.7, 1.4, 2.3, 7, 70), see Table 1.

When testing the copy number of motors on each cluster, to ensure that the results are affected by the clustering of motors rather than the total motor number, the number of clusters in the system (N_c_) was varied together with the number of kinesins on each cluster (N_k_), so that the total motor number (N_k_·N_c_) is kept constant, see Table 1.

The modified Cytosim code is publicly accessible (https://github.com/Archie-G/cytosim_MT_breaking). For parameters and configurations used in this study please see Table 1, as well as example configuration files in the GitHub repository.

### Image analysis

For Fig. 1 B, to quantify the level of KIF1C LLPS activity in cells (x-axis), the heterogeneity of KIF1C-EGFP fluorescence intensity within a cell was measured by calculating the intensity entropy:

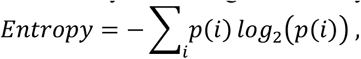

where *p(i)* is the probability of the i-th intensity level after ranking the intensity of all pixels within a selected cell (outlined manually) in an 8-bit image. This entropy value reflects the heterogeneity of KIF1C-GFP fluorescence intensity in each cell (i.e., low values indicate a homogenously diffuse KIF1C, and high values indicate a heterogenous localization of KIF1C between diffuse, small puncta, and large puncta). The number of microtubule fragments in each cell (y-axis) was counted manually.

For Fig. 6 C, 7 C and 8 E, to quantify the amount of microtubule breakage and fragmentation within a cell, we took the number of microtubule free ends and normalized it to the microtubule area in a 20 μm x 20 μm area at the periphery of a cell. Briefly, a 20 μm x 20 μm subcellular region of interest (ROI) was selected where KIF1C-EGFP condensates were localized at the edge of the cell. A normalized microtubule fragmentation value for the ROI was obtained by counting the number of microtubule free ends and dividing it by the relative microtubule area within the region. The number of microtubule free ends was marked manually, whereas the microtubule area (in μm^2^) was obtained by thresholding using Auto Local Threshold (Phansalkar method selected). Therefore, this normalized value represents the density of microtubule free ends relative to the area of microtubule.

### Quantification and statistical analysis

Statistical analyses were performed and graphs were generated using R and RStudio. The mean and standard error are described in the main text and/or the figures. The number of values examined, the experimental replicates, and the statistical tests applied are described in the figure legends.

## Acknowledgements

We thank François Nédélec for discussions and advice on the use and modification of Cytosim. We thank David Sept and Patrick DeLear for their advice and feedback on the simulations. We are grateful to Ryoma (Puck) Ohi, Morgan DeSantis, Michael Cianfrocco and members of their laboratories for helpful discussions and reagents. We are grateful to Eric Rentchler in the University of Michigan Biomedical Research Core facilities. This work was supported by grants from the National Institutes of Health to K.J.V. (R35 GM131744). S.N.S was supported by NIH T32 GM145470.

---

**Figure S1.**
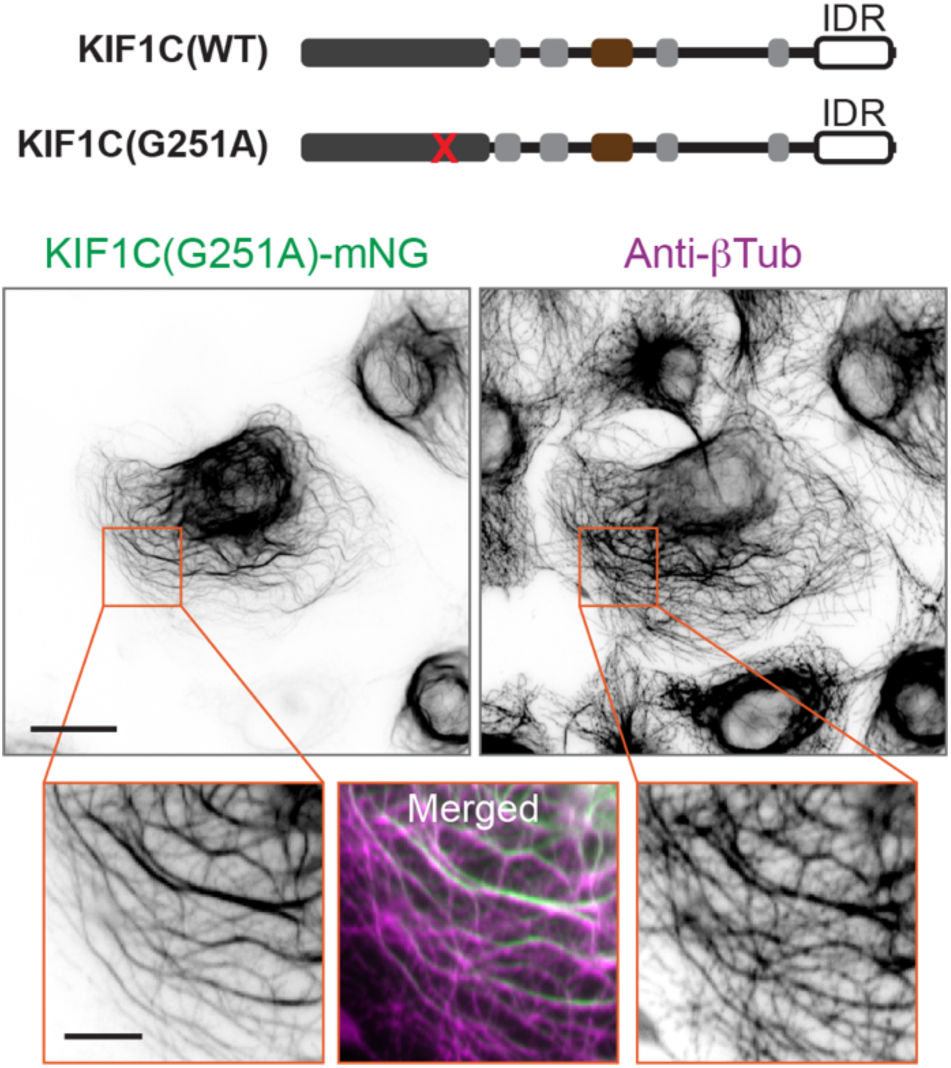
A rigor mutant of KIF1C does not induce MT fragmentation. (Top) Schematics of the domain organization of full-length KIF1C. The red X marks the location of the G251A rigor mutation. (Bottom) Representative immunofluorescence images of KIF1C(G251A)-mNG expressed in COS-7 cells and fixed and stained for microtubules with anti-βTub. Orange boxes indicate the regions shown in the magnified images. Scale bars: 20 μm for whole-cell views and 5 μm for magnified images.

**Figure S2.**
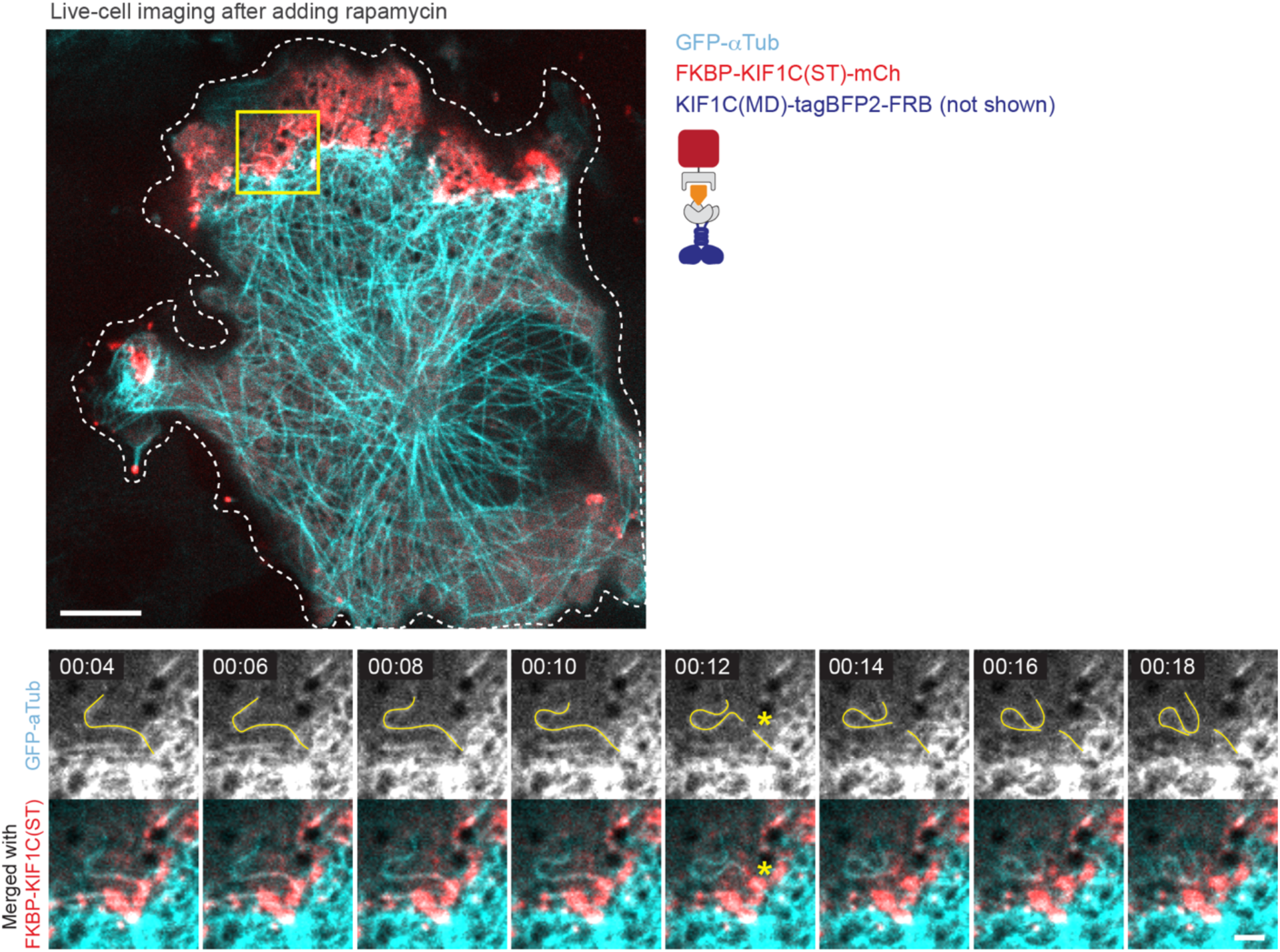
Live imaging of MT breakage by KIF1C condensates. A second representative example (with Fig. 3) of live-cell imaging of COS-7 cells expressing KIF1C(MD)-tagBFP2-FRB (imaging channel not shown), FKBP-KIF1C(ST)-mCherry (magenta), and EGFP-α-tubulin (cyan). The cells were imaged at 2 frames per second starting 10-15 min after rapamycin addition. Dashed white lines mark the cell outline, the yellow box indicates the region magnified in the time series below. Time stamps are in min:sec. The yellow asterisk marks the position of MT breaking. Scale bars: 10 μm in whole-cell view and 2 μm in magnified views.

**Figure S3.**
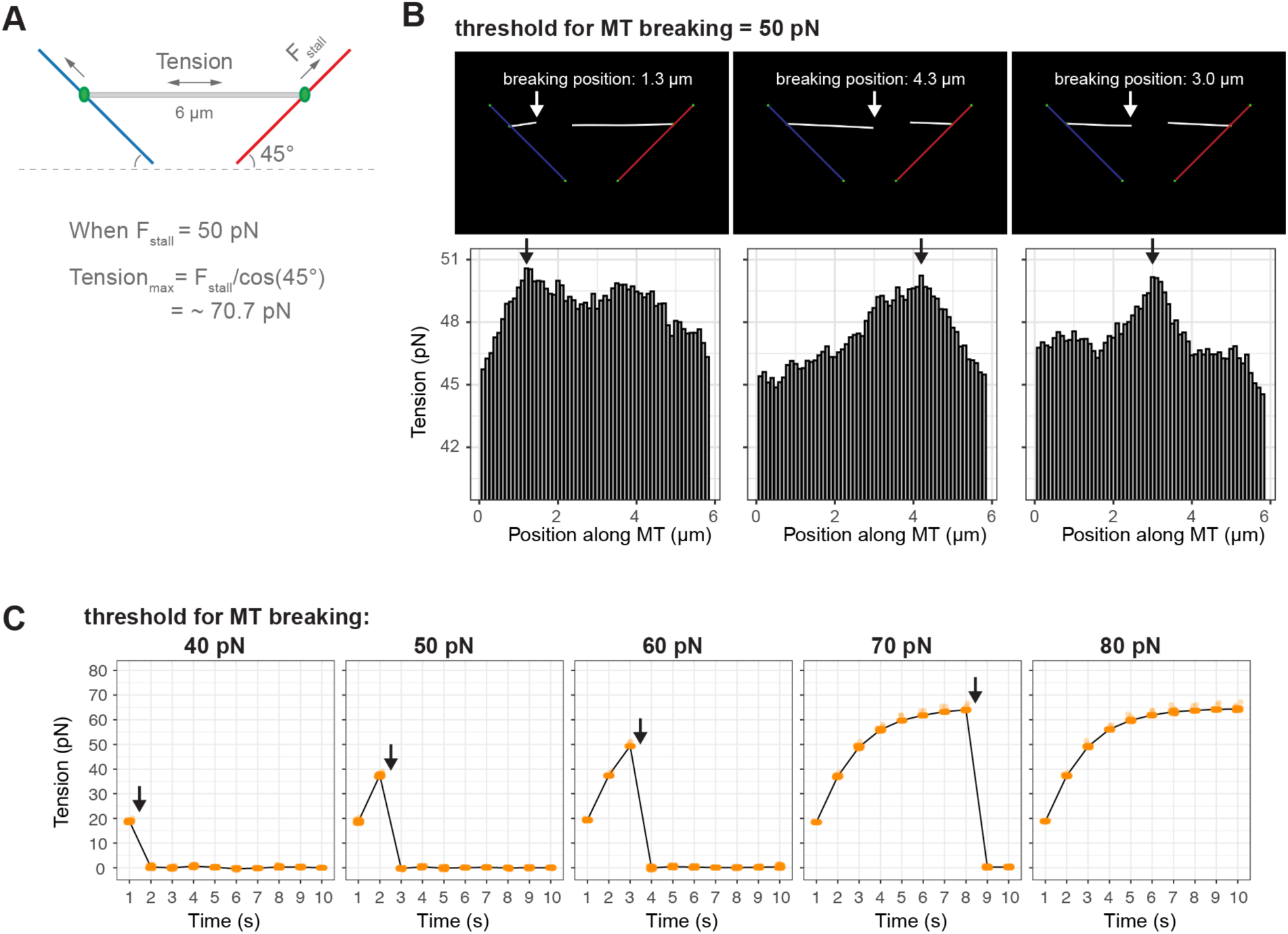
Validation of single microtubule fiber breaking behavior in Cytosim. **(A)** Schematic of simulations that apply tension to a single microtubule fiber. The gray horizontal bar represents the MT fiber (length = 6 μm, segment number = 60, length per segment = 0.1 μm). Green dots represent the artificial motors that pull the fiber in opposite directions to apply tension. Blue and red lines represent the tracks on which the artificial motors move in the direction indicated by the arrows. F_stall_: stall force of the artificial motors. Tension_max_: the highest tension that can be generated given the stall force of the artificial motors. **(B)** Three representative simulation results with F_stall_ and the threshold for microtubule breaking set to 50 pN. Top: images of simulation space after microtubule fiber breaking, with the position of the rupture event indicated by the white arrows. Bottom: graphs displaying the tension values along the microtubule fiber at the moment of rupture, with the black arrows pointing to the position of breaking. The position of highest tension on each fiber correlates with the position of breaking. **(C)** Microtubule breaking at different threshold levels with F_stall_ set at 50 pN. The x-axis shows the time-course of the simulation; the y-axis shows the tension on the microtubule fiber. The orange dots display the tension at each segment (0.1 μm per segment) of the microtubule fiber. A sudden drop of tension on the microtubule fiber (black arrows) indicates a microtubule breakage event.

**Figure S4.**
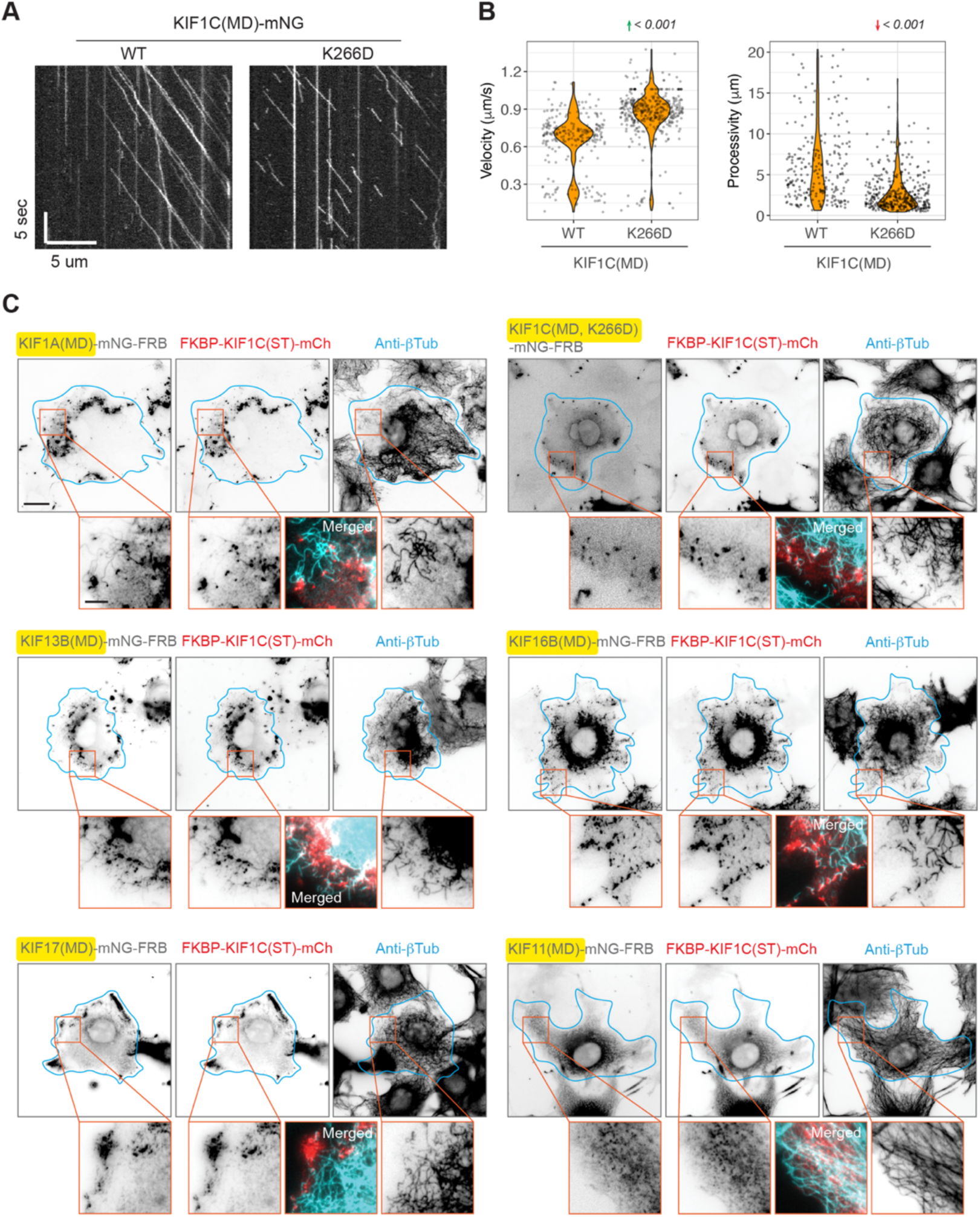
Split-kinesin assay using kinesin motor domains with different biophysical properties. (A,B) The K266D mutation results in a decrease in KIF1C processivity. **(A)** Representative kymographs from *in vitro* single-molecule motility assay using lysates of COS-7 cells expressing wild-type (WT) or mutant (K266D) KIF1C(MD)-mNG. Time is on the y-axis, and distance is on the x-axis. Scale bars are shown on the graph. **(B)** Violin plots of motor velocity and processivity (run-length) measured from single-molecule motility assays. Each dot represents one motility event. Data were collected from 2 independent experiments. p-values (unpaired t-test) are shown on top of the graphs. Red arrow indicates a decrease compared to WT; green arrow indicates an increase compared to WT. **(C)** Representative immunofluorescence images of split-kinesin assays in COS-7 cells using various kinesin motor domains assembled into multi-kinesin clusters via the KIF1C(ST) domain. Microtubule breakage is quantified in Fig. 6 C. Cyan lines mark the cell outlines, orange boxes indicate the regions shown in magnified view below. Scale bars: 20 μm for whole-cell views and 5 μm for magnified images.

**Figure S5.**
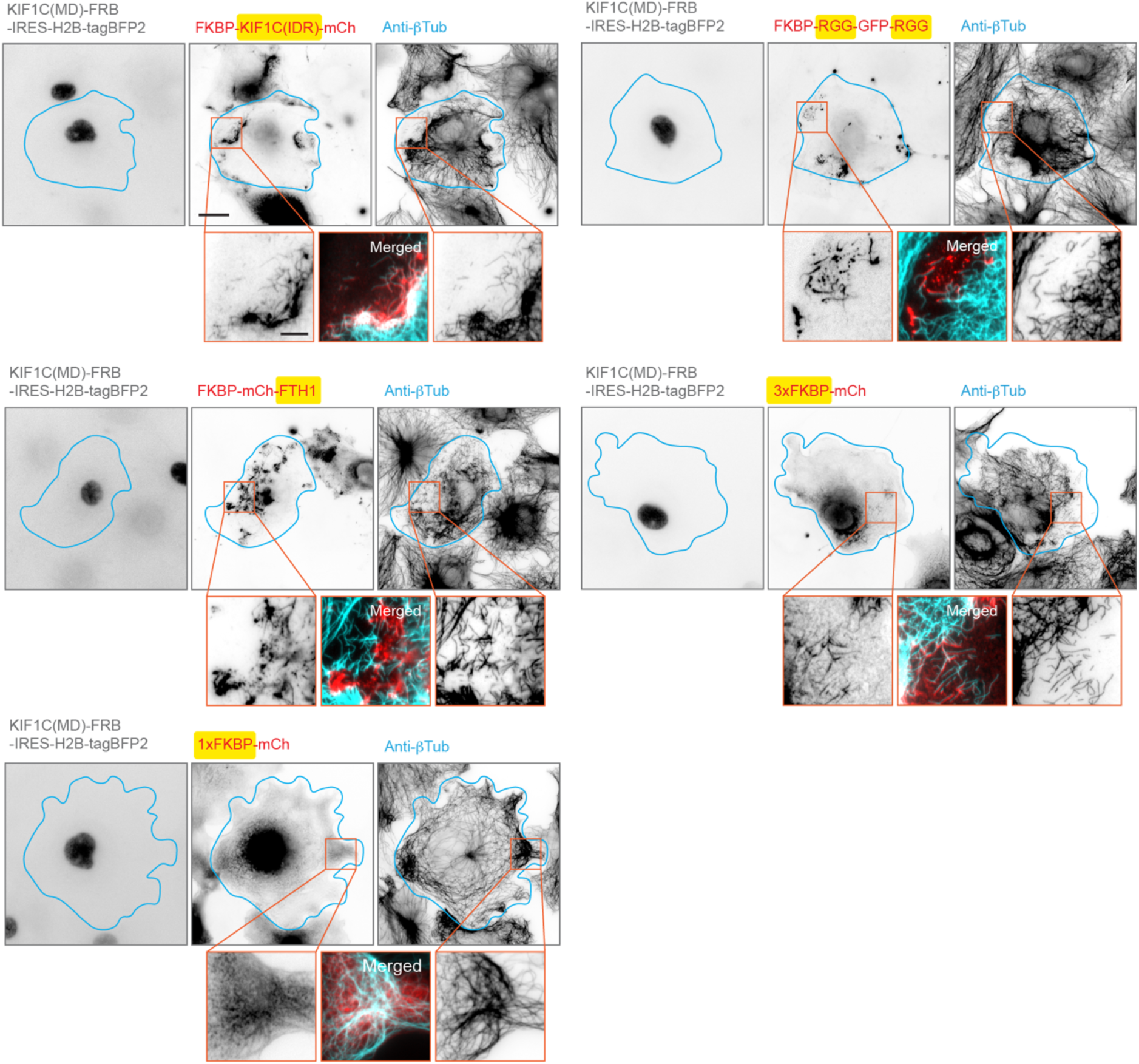
Split-kinesin assay using clustering domains with different properties. Representative immunofluorescence images of split-kinesin assays in COS-7 cells using various clustering mechanisms to generate multi-kinesin clusters of the KIF1C(MD). Microtubule breakage is quantified in Fig. 7 C. Cyan lines mark the cell outlines, orange boxes indicate the regions shown in magnified view below. Scale bars: 20 μm for whole-cell views and 5 μm for magnified images.

**Figure S6.**
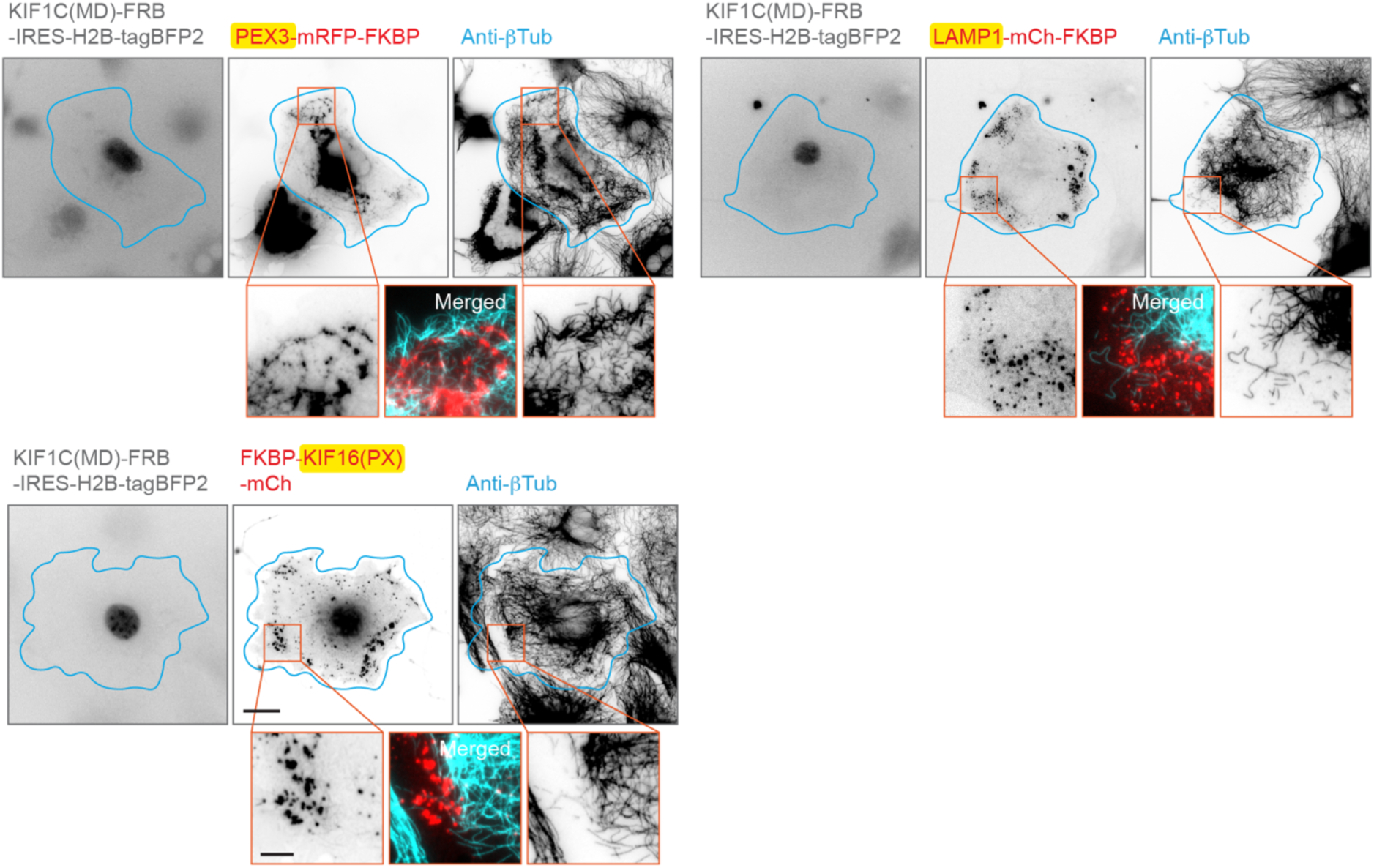
Split-kinesin assay using different motor-cargo attachment mechanisms. Representative immunofluorescence images of split-kinesin assays in COS-7 cells using various mechanisms of motor-cargo association to assemble multi-kinesin clusters of the KIF1C(MD). Microtubule breakage is quantified in Fig. 8 D. Cyan lines mark the cell outlines, orange boxes indicate the regions shown in magnified view below. Scale bars: 20 μm for whole-cell views and 5 μm for magnified images.

